# An integrative approach: using transcriptomic data and network analysis of transcriptional reprogramming in tomato response to PSTVd variants

**DOI:** 10.1101/2022.02.02.478822

**Authors:** Katia Aviña-Padilla, Octavio Zambada-Moreno, Gabriel Emilio Herrera-Oropeza, Marco A. Jimenez-Limas, Peter Abrahamian, Rosemarie W. Hammond, Maribel Hernández-Rosales

## Abstract

Viroids are minimal pathogens of angiosperms, consisting of non-coding RNAs that cause severe diseases in agronomic crops. Symptoms associated with viroid infection are linked to developmental alterations due to genetic regulation. To understand the global molecular mechanisms of host response, we implemented an omics approach to identify master transcription regulators (MTRs) and their differentially expressed targets in tomato infected with mild and severe variants of PSTVd. Our approach integrates root and leaf transcriptomic data, gene regulatory network analysis, and identification of affected biological processes. Overall, our results reveal that specific bHLH, MYB, and ERF transcription factors regulate genes involved in molecular mechanisms underlying critical signaling pathways. Functional enrichment of regulons shows that bHLH-MTRs are linked to metabolism and plant defense, while MYB-MTRs are involved in signaling regulation and hormone-related processes. Strikingly, a member of the bHLH-TF family have a potential specific role as a microprotein involved in the post-translational regulation of hormone signaling events. For the severe variant, ERF-MTRs are characteristic, while ZNF-TF, tf3a-TF, BZIP-TFs, and NAC-TF act as unique MTRs. Altogether, our results lay a foundation for further research on the PSTVd and host genome interaction, providing evidence for identifying potential key genes that influence symptom development in tomato plants.

## 1. Introduction

Viroids are plant restricted parasites that cause significant losses in agronomically important crops. They are composed of non-coding single-stranded, circular RNA (246-491nt) of low molecular weight without a protein or lipid membrane[1]. The mechanism of systemic infection consists of hijacking the machinery of the host cell by interacting with cellular factors for their replication[2]. The proposed pathogenesis mechanisms may be regulated by the direct interaction between viroid genomic RNA, and/or replication intermediates, and the host genome[3]. Environmental factors such as photoperiod, temperature, and nutrient availability are crucial for viroid infection and may determine whether host plants are asymptomatic or exhibit a wide range of symptoms from mild to the severe expression. Symptom development is an outcome of the complex plant-pathogen interaction as a result of alterations to cell processes related to viroid RNA-mediated genetic regulation.

Genetic regulation is a highly complex process that comprises concerted events by multiple layers of the interplay among DNA, RNA, and proteins. Plant response to pathogen infection conditions involves a precise reprogramming of host transcriptional activity. The increase in data availability has led to an interest in systems biological approaches in plants to characterize the molecular mechanisms and hub genes relevant to a specific tissue or disease condition. Recent advances employing next-generation sequencing have led to a better insight into the molecular biology of viroid-host interactions. Transcriptional profiling analyses have revealed that viroid infections have a global effect on plant gene expression. These studies include potato spindle tuber viroid (PSTVd) infection in tomato [4–7] and pepper [8], citrus exocortis viroid (CEVd) [9], and citrus viroid III (CVd-III) [10] infection in Etrog citron, peach latent mosaic viroid (PLMVd) infection in peach [11], hop stunt viroid (HSVd) in hop [12] and cucumber [13], and hop latent viroid (HLVd) and citrus bark cracking viroid (CBCVd) in hop [14].

Altogether, these previous studies showed alterations in host genes linked to defense, stress response, hormone signaling, DNA-binding, and RNA metabolism, among other functions [4–14].

In keeping with this, diverse studies have recently reported several DE-TFs in the tomato and pepper host responses to PSTVd infection, highlighting that in-depth studies approaching the specific role of each of them must be considered to clarify their link to disease phenotypes[7–8]. Taking advantage of data expansion, we used publicly available transcriptome profiles from microarray technology of time-course studies of mild (M) and severe (S23) PSTVd infections in leaf and root samples derived from tomato plants [7–15]. Results from these studies highlighted the differences in gene expression profiles depending on the infection stage and the PSTVd strain. In our study, we focused on performing network analysis using integration of these data, since it represents the most comprehensive available concerning transcriptomic analysis comparing mild and severe pospiviroid infections. In this context, computational strategies using transcriptomics data integration for network inference to study complex diseases have become evident [16–20]. Data integration allows to include more samples providing increasing information. In other words, the consistency of co-expression networks increases as the sample size increases [20–23]. The aim of this work is to explore the mechanisms regulating gene expression linked to host symptoms, at the transcriptional and post-translational levels.

Transcription factors (TFs) are central regulators of gene expression, and their role relies upon their capacity to specifically interact with DNA sequences and other proteins as part of transcriptional complexes modulating key features of organismal biology, including cell differentiation, tissue and organ development, responses to hormones and environmental factors, metabolic networks, and disease resistance, among others. TFs regulate the transcription of downstream targets with the interaction of co-regulators known as regulons [24]. Tomato (*Solanum lycopersicum* L.) is an agronomically important crop and has been used as a model for basic research in plant development [25]. This crop is one of the most widely grown greenhouse vegetables in the world [26]. In addition, it is one of the most profitable and widely consumed by its various products. During 2018, 243,888,041 tons were harvested [27]. Diseases caused by viroids cause serious economic losses in this host [28–30]. A high-quality genome sequence for domesticated tomato has been assembled by the International Tomato Genome Sequencing Project and more than 34,000 proteins have been predicted, with functional descriptions assigned to 29,532 genes (http://solgenomics.net/organism/Solanum_lycopersicum/genome). The tomato genome contains 1845 genes that code for transcription factors (TFs), [31]. The transition from vegetative growth to reproductive development requires gene network coordination, where TFs act as essential regulators of organ morphogenesis. Our study focused on those TFs that could play a key role in the expression of genes involved in biological processes linked to viroid-induced symptoms. For that purpose, gene regulatory networks are proposed to infer an interaction mechanism. We used 53 expression profiles for three different PSTVd infection stages of mild (M) and severe (S23) variants. In this work, we identified Master Transcriptional Regulators (MTRs), particularly in the (basic helix-loop-helix) bHLH, MYB, and (ethylene-response factors) ERF superfamilies, that have been shown to have a relevant role in viroid infection [7–9]. In plants, these TFs regulate growth, development, and stress response [32–41].

Moreover, there is recent evidence for the formation of highly regulated protein complexes through protein-protein interaction domains, the disruption of which can have serious consequences for cell function. In this context, the formation of dimers and multimers can be disturbed by proteins known as microproteins (miPs),[42–43]. We have recently reported the role of miPs of the conserved bHLH superfamily, which has evolved in both the animal and plant kingdoms, [44]. miPs regulate multidomain proteins at the post-translational level. They are analogous to microRNAs, having the capability to heterodimerize with their targets, causing dominant and negative effects. In plants, bHLH-TFs have recently been characterized as miPs involved in cell elongation, brassinosteroid events, and plant development [45–46].

Our study integrates network models of genes affected during viroid infection in tomato, proposing mechanisms of host gene regulation involving MTRs and targets under their regulation, as well as potential specific miPs. Overall, our results delineate the highly specific and pivotal role that bHLH, MYB, and ERF MTRs play in coordinating gene regulatory networks in PSTVd-infected tomato hosts. Strikingly, our analysis also highlights specificity at a post-translational regulation level involving bHLH miPcandidates linked to auxin-responsive factors, jasmonate, and brassinosteroid pathways, with a distinctive signature for the tomato response to infection with the severe strain of PSTVd.

The replication of pospiviroid species takes place in the nucleus and regulates the expression of key genes by a still unknown mechanism that involves the transcriptional reprogramming by host-encoded biomolecules. Identifying the key regulators, the interacting miPs, as well as the downstream differentially expressed genes will contribute to unraveling the pathogenesis mechanism responsible for the characteristic phenotypes shown in the susceptible infected plants.

## 2. Results

### 2.1 Gene Regulatory Networks (GRNs) identify specific bHLH, ERF, and MYB TFs that act as master transcriptional regulators in the tomato response to PSTVd infection

We used integrated transcriptome microarray data for time-course analyses under three experimental conditions: Healthy Control (C.); PSTVd-Mild (M) and PSTVd-Severe (S23) infections in roots (GSE111736) and leaf (GSE106912) tissue. Gene regulatory networks (GRNs) were analyzed for the three treatments. Then, we performed Master Regulator Analysis (MRA), using a network model derived from the largest available viroid tomato microarray datasets from GEO [12,13]. The analysis allowed us to generate a robust readout not only of the changes of the transcript of each protein-encoding gene in the interactome but also to study its proximal functional network, [47].

The global GRN of the integrated samples is depicted in **Figure S1a**. Altogether, the GRN analysis showed that the S23 strain interactomes are denser than those characteristics of the control and mild conditions, highlighting the relevance of the activation of plant defense mechanisms as a response to the viroid virulence. The MRA results showed that out of the 1845 transcription factors in the tomato genome, eighty-seven are MTRs modulating biological processes among the three different conditions, **Figure S1b**.

In the C. vs M. comparison, we identified seven significant MTRs, while, for the C. vs. S23 condition, we found fifty-nine significant MTRs. On the other hand, when comparing M. vs. S23 the results indicated twenty-one significant MTRs, see **Table S1**. As could be expected for the significant differences in the plant phenotypes underlying the response to the aggressiveness of the viroid variant, the highest number of MTRs (more than 60% of the significant MTRs) is found in the C. vs. S23 comparison.

The most significant MTRs identified in the C. vs S23 condition are, pivotal TFs such as TCP *(Solyc01g008230.2); Solyc06g070900.2,* ZINC FINGER TF15 *(Solyc01g110490.2),* MADS-box *(Solyc01g087990.2), BZIP(Solyc01g111580.2)*, and GRAS *(Solyc05g053090.1)* members. Notwithstanding, among the significant identified MTRs, TFs of the bHLH, MYB, and ERF families stand out in the MRA analysis. Notably, in the C. vs S23 comparison five members of the bHLH family SlbHLH0ll *(Solyc01g111130.2)*, bHLH130-like *(Solyc12g100140.1),* SlbHLH022 *(Solyc03g097820.1);* GBOF-*1(Solyc06g072520.1)*, and bhlh92 isoform *(Solyc09g083360.2)* are included. The following three MYB TFs, MYBlRl *(Solyc04g005100.2), MYB16(Solyc02g088190.2)*, kua1 isoform *x1(Solyc08g078340.2);* as well as seven ERFs: ERF-1a *(Solyc05g051200.1)*, ERF_C_5 *(Solyc02g077370.1)*, AP2/EREBP TF1 *(Solyc02g093130.1), RAP2-12(Solyc12g049560.1),* PTI6 *(Solyc06g082590.1),* TSRF1 *(Solyc09g089930.1),* and ERF_A_2 *(Solyc03g093610.1)* were identified.

While in the M vs S comparison two bHLH-MTRs were found. bHLH0lO a cryptochrome-interacting basic-helix-loop-helix *(Solyc01g109700.2)* and MYC2 a jasmonate related *TF(Solyc08g076930.1)*. While two MYB-TFs, kua1 isoform *x1(Solyc08g078340.2)*, PHR1-LIKE 1*(Solyc05g055940.2)*, and two ERF TFs AP2/EREBP TF1 *(Solyc02g093130.1)*, ERF_C_5 *(Solyc02g077370.1)* were identified. In the C vs M comparison only a MYB-TF *Solyc03g098320.2* is participating as an MTR. In total, seven bHLH, seven ERF and five MYB unique members were identified as MTRs in the analyzed conditions. We selected and focused on the members of those families to delve insight into their roles based on their representation and high specificity in the different analyzed treatment conditions.

### 2.2 Gene communities in the bHLH, ERF, and MYB interactomes highlight their functional significance in specific and shared biological processes

The identification of communities, or modules, is a common operation in the analysis of large biological networks. In order to determine mechanisms of regulation in our global GRN, we used the Louvain-Method that allows us to identify gene communities that could be participating together in particular biological processes, [48]. For that purpose, we split our global GRN into subnetworks of the **bHLH**, MYB, and ERF MTRs interactomes with their regulons.

Our analysis revealed that in the interactome of the bHLH MTRs four biological communities are found. See **Appendix A** for communities description, and **Appendix B** for all the functional enrichment analysis performed in this study. The first contains 2040 genes enriched in GO:0015979 photosynthesis (*padj-value* = 0.0000015). A second community containing 1439 genes functionally enriched in KEGG:00660-C5-Branched dibasic acid metabolism and (*padj.-value* = 0.002571), and KEGG:00290-Valine, leucine, and isoleucine biosynthesis pathways were identified (*padj-vahte*=0.0276). While, the third contains 877 genes linked to the KEGG:03010-Ribosome pathway, GO:0042254 ribosome biogenesis, and anthocyanin biosynthesis, (*padj.-value* >0.000003-0.01352). Finally, in the fourth community that includes 870 genes, an enrichment in DNA-binding transcription factor activity and specific mRNA binding was determined, (*padj-value*= 3.969E-2).

Regarding the ERF subnetwork, ten gene communities with biological significance were found. This is the interactome with the higher number of communities found participating in multiple biological procceses. The community with the highest number of genes has 578 enriched in the GO:0045272-plasma membrane respiratory chain complex *(padj.-value*= 0.00254). Meanwhile, a second community of 280 genes that are enriched in protein self-association, aromatic amino acid family biosynthetic process, and regulation of cellular biosynthetic process was identified *(padj-value* 4.507E-2 >3.343E-2). The third includes 268 genes enriched in the KEGG:04626-plant-pathogen interaction pathway *(padj.-vahte=0.00121)*. While a community of 247 genes is enriched in the KEGG:00750-Vitamin B6 metabolism *pathway(padj.-value* =0.03634), and 201 genes protein heterodimerization activity, serine O-acyltransferase activity, and, nucleosome organization and packaging *(padj.-value* =0.03634). Two communities of 158 genes were found and are related to the KEGG:00660-C5-Branched dibasic acid metabolism *(padj.-vahte=0.0399)* and ribosomal large subunit biogenesis. Those pathways are shared with the communities found in the bHLH TF interactome. In addition, 95 genes participate in a community involved in the regulation of the cell cycle *(padj-value*= 9.041E-3), 81 genes enriched in regulation of cell wall pectin metabolic process, self proteolysis, positive regulation of response to stimulus, and positive regulation of MAP kinase activity *(padj-value*= 4.659E-2>2.800E-2), while the last community comprises 71 genes that are enriched in the KEGG:00942-Anthocyanin biosynthesis pathway *(padj.-value* =0.01998).

Finally, when performing the operation for the MYB TFs interactome, the results revealed five gene communities participating in additional processes. For instance, 1852 genes in GO:0015979 photosynthesis (*p-value* =2.8596E-10); 1804 genes in cellular divalent inorganic cation homeostasis rRNA binding GO:0019843 (*p-value* =0.01183) 1715 genes in the following biological processes: GO:0006950 response to stress, GO:0006952 defense response and the KEGG:04626 Plant-pathogen interaction pathway (*p-value* 0.0024>0.0367); while 875 genes were found to be related to the GO:0031365N-terminal protein amino acid modification (*padj-value*= 0.0327) The presence of a small community of 12 genes participating in amino acid and carbixylic acid trasmembrane transport (*padj-value*= 6.839E-3) was also found.

In summary, our results revealed that bHLH, ERF, and MYB TFs regulate gene expression in communities associated with unique biological processes such as the valine, leucine, and isoleucine biosynthesis, vitamin B6 metabolism, and anthocyanin biosynthesis, amino acid and other molecule; transmembrane transport, and KEGG pathways, as well as participating together in the modulation of conserved crucial molecular processes like the C5-Branched dibasic acid metabolism, ribosome, photosynthesis, plant-pathogen interaction and defense response in the PSTVd variants infection.

### 2.3 bHLH MTRs are specific modulators of genes involved in metabolism, ribosome, light, and plant defense processes

When analyzing the interactome containing thirty-two co-expressed bHLH TFs and their regulons, a total of 4411 nodes and 7424 interactions were obtained. For the C. vs. S23 comparison, five unique members of this family were classified as MTRs, **Figure 1a**.

**Figure 1.**
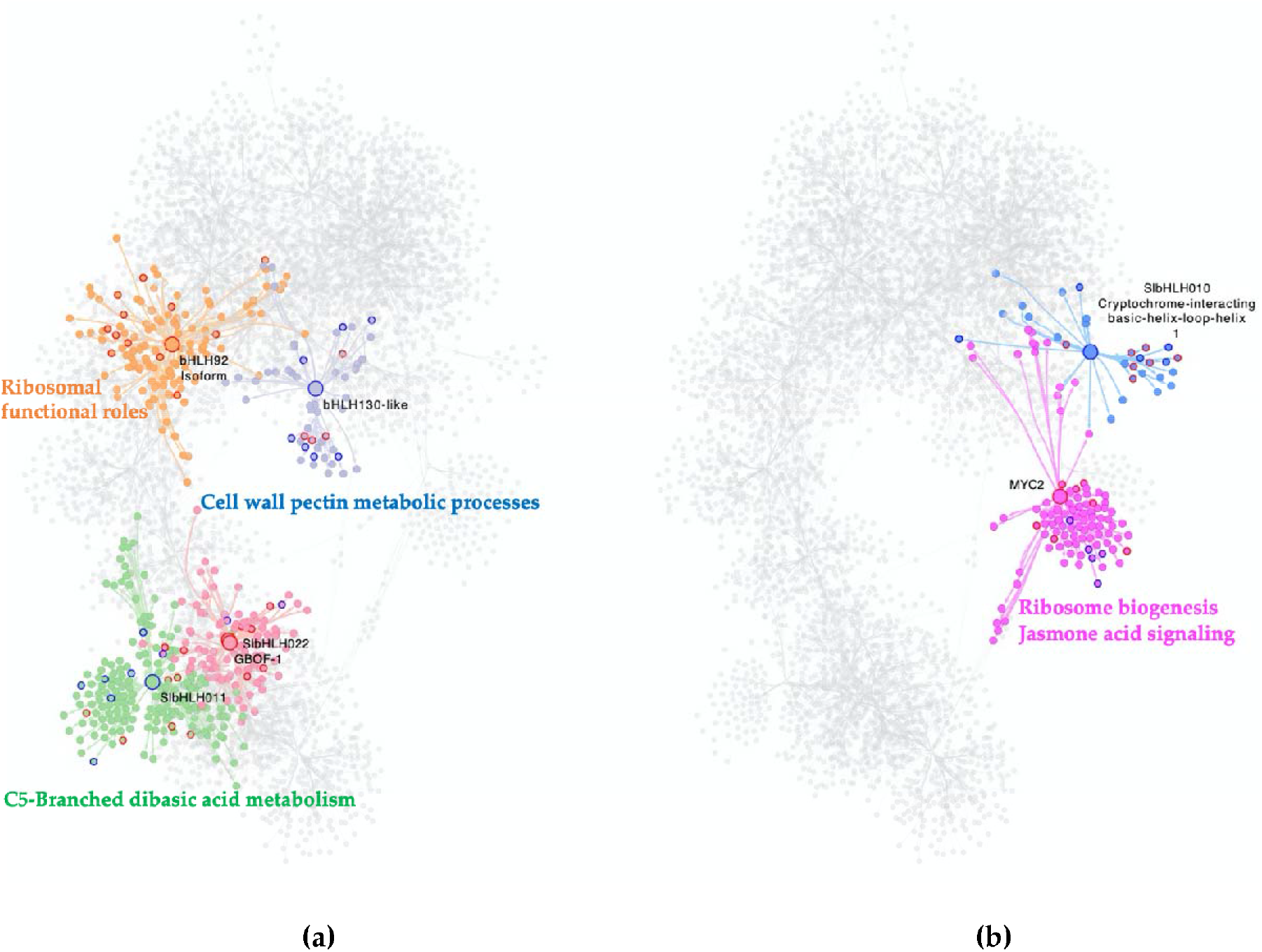
Interactome of the bHLH MTRs in the experimental conditions comparisons: **(a)** C vs S23 condition, (b): M vs S23 condition.. Each node represents a gene, the edges between nodes represent regulatory interactions between genes. The big highlighted nodes with a specific color represent bHLH-MTRs the coloured perimeter of each MTR represents their NES value, while nodes of the same color surrounding these represent target genes regulated by an MTR, the most differentially expressed genes in each regulon have coloured perimeters, blue for down regulated, red for up regulated. Enriched biological processes of each regulon are presented in the same color as the regulon. In the comparison M vs S23: bHLH-MTRs are MYC2 (pink), SlbHLH0lO (blue). Meanwhile in the comparison C vs S23 bHLH-MTRs are bHLH130-like (blue), bHLH92 (orange), SlbHLH011 (green), SlbHLH022 (cherry red), GBOF-1 (cherry red).

The MRA results showed induction of their regulons SlbHLH022 *(Solyc03g097820.1);* GBOF-1(*Solyc06g072520.1)* with unknown functions; and bhlh92 isoform *(Solyc09g083360.2)* enriched in ribosome and ribonucleoprotein complex biogenesis functional roles, **Table S1; Figure 1a**. In contrast, the inhibition of regulons under the modulation of SlbHLH0ll *(Solyc01g111130.2)*, linked to C5-Branched dibasic acid metabolism, and bHLH130-like *(Solyc12g100140.1)*, related to cell wall pectin metabolic processes, was identified in **Table S1; Figure 1a**. In the comparison of the M. vs. S23, the relevance of two bHLH TFs was revealed in the MRA analysis.

The induction of the regulon of MYC2 *(Solyc08g076930.1)* is related to ribosome biogenesis and jasmonic acid signaling, while the repression of the bHLH0lO a cryptochrome-interacting basic-helix-loop-helix *(Solyc01g109700.2)* has no functional enrichment, however this gene is related to triggering flowering in response to blue light,[49], **Figure 1b**.

Strikingly, none of the bHLH-identified MTRs are shared between the C, M, and S23 treatment conditions. Overall these results suggest that bHLH reprogramming of regulatory networks could be specific to PSTVd variant infection in its tomato host.

### 2.4 MYB MTRs are associated with gene regulation in cell division, signaling, and mRNA transcription

An interactome network containing thirty-nine MYB TFs from the GRN was generated to give deep insight and a better understanding of the role of this TF family in viroid infection and symptom development. The MYB interactome is composed of 4380 nodes with 7312 interactions among them.

Our MRA results show that for the C. vs. S23 comparison a total of three MYB-TFs are MTRs **Figure 2a**. Being the following two modulating a lower expression of their regulons: *MYB16(Solyc02g088190.2)* related to kinase inhibitor activity, sequence-specific mRNA binding, and kua1 isoform *x1(Solyc08g078340.2)* which is enriched in the regulation of mitotic sister chromatid separation, and rRNA metabolic process.Meanwhile, upregulation of downstream gene expression of MYBlRl *(Solyc04g005100.2)* regulon is implicated in anatomical structure development, meiosis I cell cycle process, meiotic nuclear division was identified.

**Figure 2.**
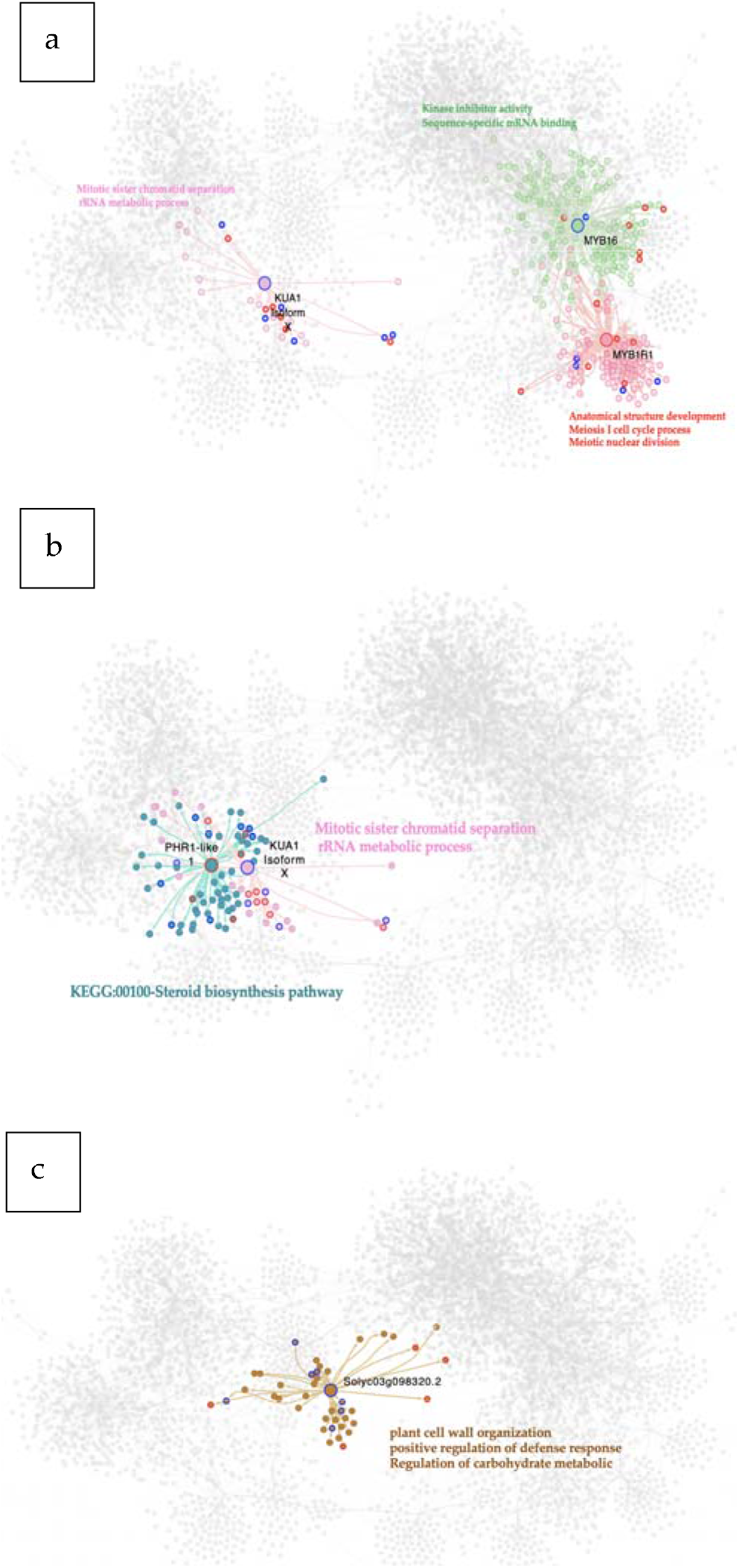
Interactome of the MYB MTRs in the experimental conditions comparisons: (a) C vs S23 condition, (b) M vs S23 condition, (c) C vs M condition. Each node represents a gene, the edges between nodes represent regulatory interactions between genes. The big highlighted nodes with a specific color represent MYB-MTRs, while nodes of the same color surrounding these represent target genes regulated by an MTR, the most differentially expressed genes in each regulon have colored perimeters, blue for down regulated, red for up regulated. Enriched biological processes of each regulon are presented in the same color as the regulon. In the comparison C vs S23 MYB-MTRs are, KUA1 (pink), MYB16 (green), and MYB1R1 (cherry red). Meanwhile in the comparison M vs S23 MYB-MTRs are PHR1 like (aqua) and KUA1 (pink). For the M vs C comparison Solyc03g098320.2 (brown) is a MYB-MTR.

For the M. vs. S23 condition, two MYB TFs act as MTRs. The gene that encodes a kua1 isoform x1*(Solyc08g078340.2)* and represses the expression of its target genes that are involved in the regulation of the mitotic cell cycle, while PHR1-LIKE 1 *(Solyc05g055940.2)* regulon shows an increased expression and is associated with the KEGG:00100-Steroid biosynthesis pathway, **Figure 2b**.

Notably, a member of this family was the only MTR found in the C. vs M. strain comparison, **Figure 2c**. This MTR gene is *Solyc03g098320.2* which represses the expression of its downstream targets, which are enriched in plant cell wall organization, positive regulation of defense response, and regulation of carbohydrate metabolic biological processes.

### 2.5 ERFs-MTRs induce the expression of their regulons as a characteristic of the tomato response to the severe PSTVd infection

As for the other TF families of interest, a subnetwork of the interactome of ERF-TFs contained in the expression data was built. This network contains 2475 nodes and 4075 interactions, being the smallest one of the three TF interactome subnetworks.

The C. vs. S23 condition analysis determined seven genes of this family as MTRs. Notably, the ERF-MTRs are abundant in the tomato response to the severe PSTVd variant. These results are in agreement with the previous PSTVd-transcriptomic studies that indicates the expression of genes encoding ERFs, Pti4, Pti6, and dehydration-responsive element-binding proteins was altered, with most being up-regulated, [7,15].The majority of the ERF-MTRs (5/7) induce the expression of their regulons ERF-1a *(Solyc05g051200.1);* ERF_C_5 *(Solyc02g077370.1);* PTI6 *(Solyc06g082590.1);* TSRF1 *(Solyc09g089930.1);* ERF_A_2 *(Solyc03g093610.1)*, while two repress the expression of their target genes AP2/EREBP TF1 *(Solyc02g093130.1); RAP2-12(Solyc12g049560.1)*, **Figure 3a**.

**Figure 3.**
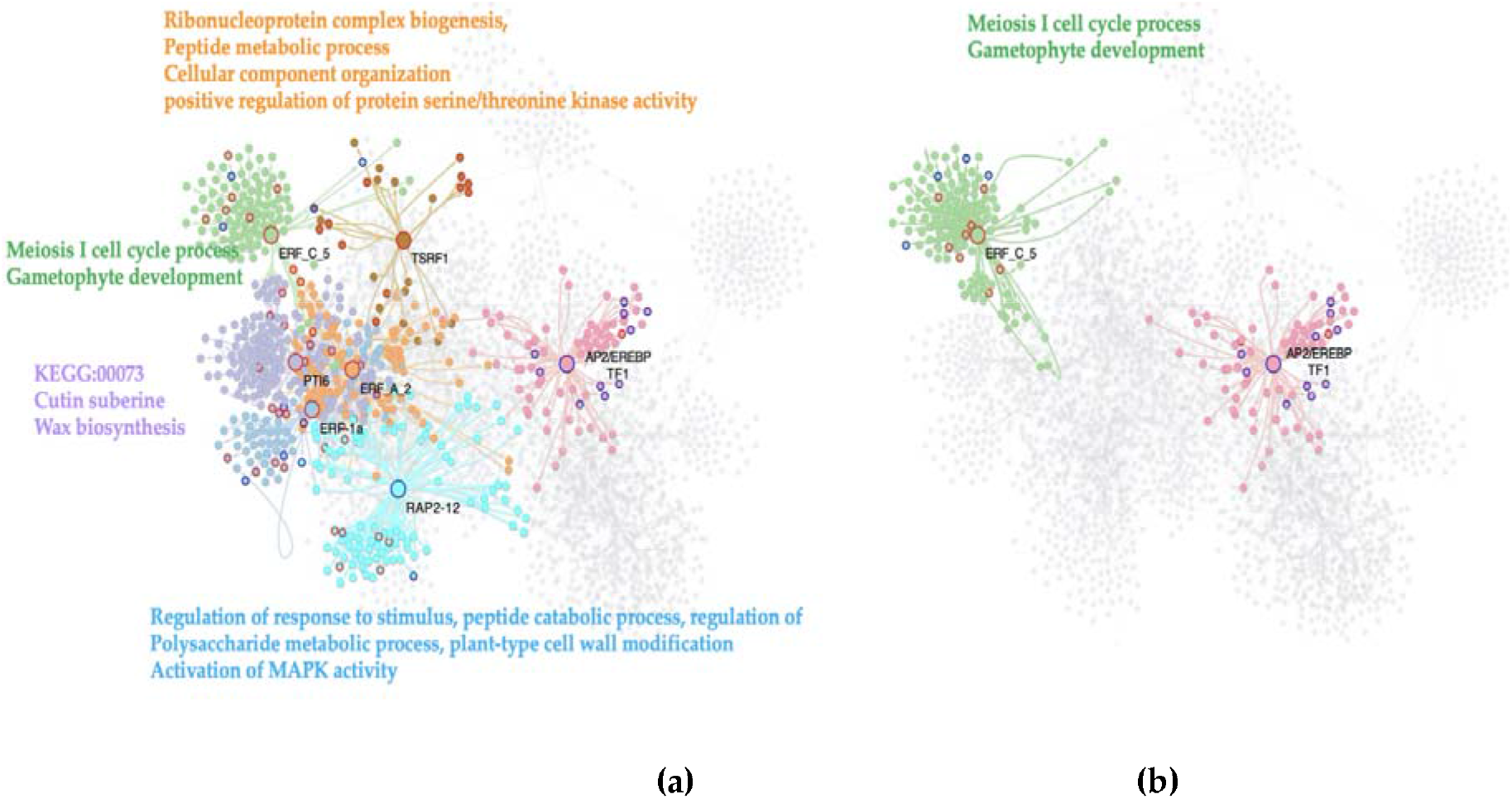
Interactome of the ERFs MTRs in the experimental conditions comparisons. (a) C vs S23 condition, (b): M vs S23 condition. Each node represents a gene, the edges between nodes represent regulatory interactions between genes. The big highlighted nodes with a specific color represent MYB-MTRs, while nodes of the same color surrounding these represent target genes regulated by an MTR, the most differentially expressed genes in each regulon have coloured perimeters, blue for down regulated, red for up regulated. Enriched biological processes of each regulon are presented in the same color as the regulon. In the comparison C vs S23 ERF-MTRs are ERF_A_2 (orange), AP2/EREBP TF1 (pink), ERF_C_5 (green), PTI6 (purple), RAP2-12 (light blue), ERF-1a (blue) and TSRl (brown). Meanwhile in the comparison M vs S23 ERF-MTRs are AP2/EREBP TF1 (pink), ERF_C_5 (green).

The induced biological processes in this comparison are resistance to fungi by ERF-1a (*Solyc05g051200.1*), which also is a key integrator of ethylene and jasmonate signals in the regulation of ethylene/jasmonate-dependent defenses, same as TSRF1 (Solyc09g089930.1). In keeping with this, the meiosis I cell cycle process, and gametophyte development are enriched terms in the regulon induced by the ERF_C_5-TF (*Solyc02g077370.1*); as well as the KEGG:00073- Cutin, suberine and wax biosynthesis synaptonemal complex organization linked to the PTI6 (*Solyc06g082590.1*). Other enriched biological procceses found in the upregulated regulos are the ribonucleoprotein complex biogenesis, peptide metabolic process, cellular component organization or biogenesis, GO:0071902 positive regulation of protein serine/threonine kinase activity by the ERF_A_2-MTR (*Solyc03g093610.1*). Alternatively, the downregulated pathways linked to ERF-MTRs in our comparison, are regulation of gene expression by stress factors and by components of stress signal transduction pathways by AP2/EREBP TF1 (*Solyc02g093130.1*), and regulation of response to stimulus, peptide catabolic process, regulation of polysaccharide metabolic process, plant-type cell wall modification, as well as, activation of MAPK activity RAP2-12(*Solyc12g049560.1*).

When analyzing the M. vs S23 comparison, two genes were selected as MTRs for the transition among phenotypes, AP2/EREBP TF1 *(Solyc02g093130.1)* it has been predicted as a transcriptional activator that binds to the GCC-box pathogenesis-related promoter element. This MTR may be involved in the regulation of gene expression by stress factors and by components of stress signal transduction pathways. In our analysis it reduced the expression of its regulon, while ERF_C_5 *(Solyc02g077370.1)* increased the expression of genes enriched in meiotic cell cycle processes, **Figure 3b**.

### 2.6 The most relevant specific MTRs for severe PSTVd variant are coupled in a negative transcriptional reprogramming

In order to identify the MTRs that are unique for the PSTVd-S23 strain and that could be correlating with the symptoms displayed by this variant, we performed an MRA and selected those that were not present for the other conditions. Our results show that 42 TFs are acting as MTRs specifically for the severe condition, **Table S2**. Among those ~30%(12/42) of the unique MTRs are TFs belonging to the families of interest. For instance, five members of the ERP-family ERF-1a*(Solyc05g051200.1);* RAP2-12*(Solyc12g049560.1);* PTI6 *(Solyc06g082590.1);TSRF1 (Solyc09g089930.1);* ERF_A_2 *(Solyc03g093610.1);* five BHLH-MTRs SlbHLH0ll *(Solyc01g111130.2);* bHLH130-like *(Solyc12g100140.1);* SlbHLH022 *(Solyc03g097820.1); GBOF-1(Solyc06g072520.1);* bhlh92 isoform *(Solyc09g083360.2)*, as well as, two MYB-type TFs, MYBlRl *(Solyc04g005100.2)*, and *MYB16(Solyc02g088190.2)*. The other remaining 70% of MTRs included diverse TFs members of multiple families including ZF-HD TFs, BZIP-TFs, NAC-TF among others, **Table S2**. We identified those that are listed in the top ten order according to their NES values and determined their predicted functional relevance.

Interestingly, in the top ten unique MTRs for this condition, TFs are mainly acting by repressing the expression of their regulons, **Figure 4**. For instance, two of them, *Solyc10g077110.1* (NES= −2.9729 *p-value*= 0.003) a tf3a transcription factor TFIIIA that plays a critical role in regulating the transcription of the 55 ribosomal RNA genes by RNA polymerase III, and BZIP23 a BZIP-family Solyc01g111580.2 (NES= −2.9667 *p-value*= 0.003) participates in the regulation of the KEGG:00660 C5-Branched dibasic acid metabolism pathway. This particular TFIIIA has been reported to be down-regulated in PSTVd-mild and S23, variants. Based on its *in vitro* binding capacity in *Arabidopsis thaliana* experimental tests, it is hypothesized that it could be acting as a bridge between the viroid template and the DNA polymerase II in the viroid-derived RNA replication, [38]. PSTVd replicates in the host cell nuclei by an asymmetric rolling-circle mechanism, where the DNA-dependent RNA polymerase forms a complex with the TFIIIA to adopt the viroid genome as a template, [50].

**Figure 4.**
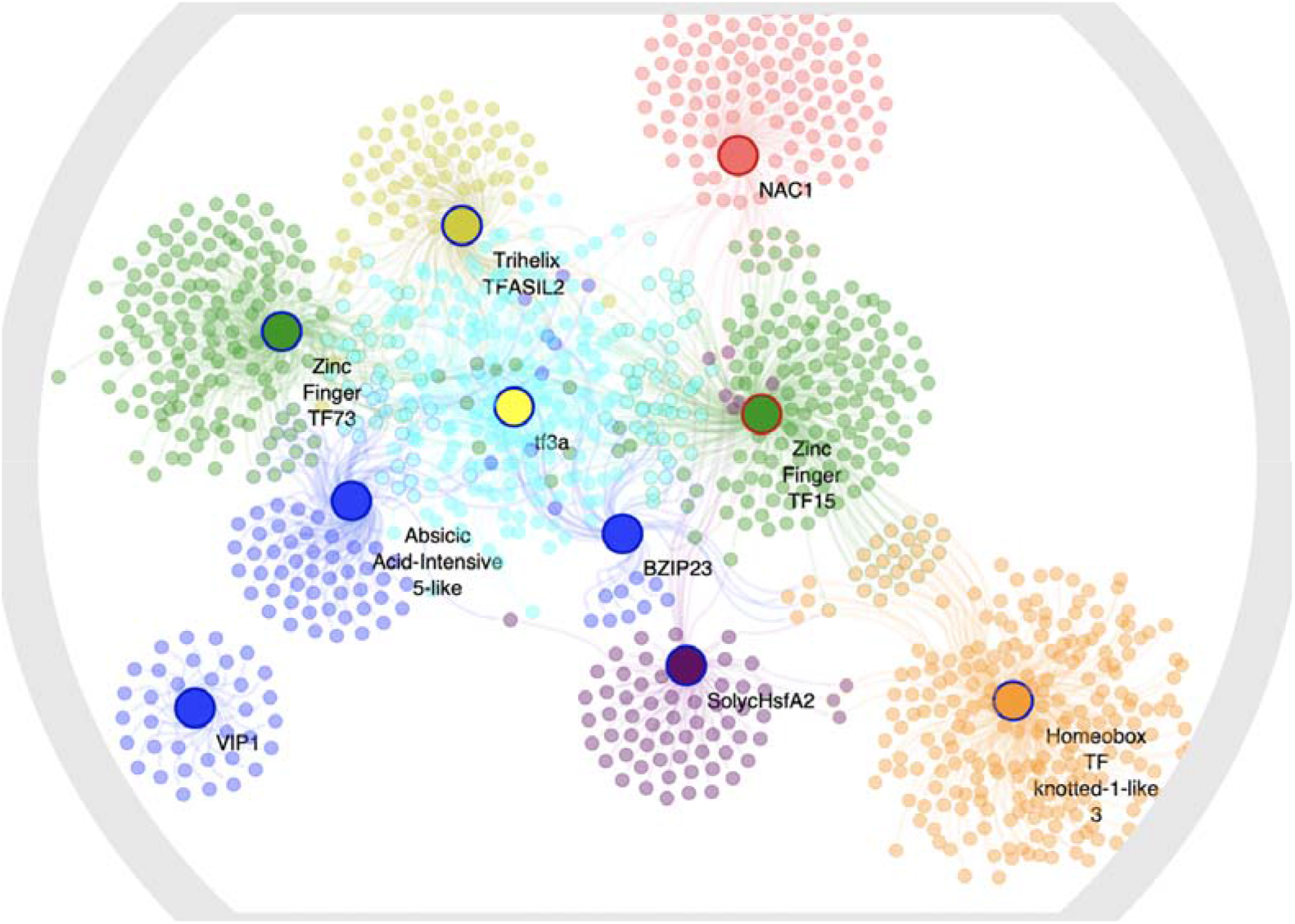
lnteractome of the top unique MTRs in the severe comparison. Each node represents a gene, the edges between nodes represent interactions between genes (regulation). Each MTR is represented by a large node, while the surrounding medium-sized nodes of the same color represent genes under its regulation. The colored perimeters in blue indicate down regulated, while in red depicts up regulation. Regulons and MTRs with the same color represent member of TF families: blue bZIP members family, while green belong to the ZF-HD family.

Notably, the presence of the BZIP-family stands out at the top of the list. Those members are *Solyc01g111580.2* (NES= −2.9667 *p-value*= 0.003) which participates in the KEGG:00660 C5-Branched dibasic acid metabolism pathway, VIP1 (*Solyc06g060490.2*) (NES= 2.8289 *p-value*= 0.004) inducing chromatin assembly, DNA conformation change, nucleosome organization, protein-DNA complex assembly, and *Solyc10g08g062960.2* (NES= −2.5241 *p-value*= 0.01) repressing cellular aldehyde metabolic process, response to desiccation, and the glyoxylate metabolic process.

Moreover, a heat shock MTR, SolycHsfA2 *Solyc08g062960.2* (NES= −2.5408 *p-value*= 0.01) which down regulates kinase inhibitor activity, and tRNA threonylcarbamoyladenosine modification in the EKC/KEOPS complex were identified. Additionally,Homeobox TF knotted-1-like 3 *(Solyc08g041820.2)* a Homeobox-MTR (NES= −2.4990 *p-value*= 0.001) downregulating sequence-specific mRNA binding, ribose-5-phosphate isomerase activity molecular functions, while NAC1 *(Solyc04g009440.2)* a NAC-MTR (NES= 2.5223 *p-value*= 0.001) inducing its regulon expression were determined, **Figure 4**.

### 2.7 A bHLH MTR has a potential specific role as miP in the post-translational regulation of hormone signaling events during severe infection

Using the miPfinder tool we obtained a list of 39 bHLH-TFs in the tomato genome as potential miPs. Out of the 39 predicted bHLH-miPs, we found that the following eight are implicit in the GRN: SlbHLH143 *(Solyc07g064040.2)*, SlbHLH14 *(Solyc02g079970.2)*, SlbHLH152 *(Solyc10g006510.2)*, SlbHLH29 *(Solyc04g006990.2), Solyc05g007210.2, Solyc03g113560.2*, SlbHLH135 *(Solyc06g050840.2)*, and the bhlh92 isoform *(Solyc09g083360.2)*. When analyzing their functional role at a protein-protein interaction level, we determined that four of these biomolecules have been previously associated with diverse plant hormones at a protein-protein interaction level, such as auxin-responsive factors *(Solyc04g006990.2, Solyc03g113560.2)*, brassinosteroids *(Solyc05g007210.2)*, as well as, jasmonate and cytokinins (bhlh92 isoform/ *Solyc09g083360.2)*, **Figure 5a**.

**Figure 5.**
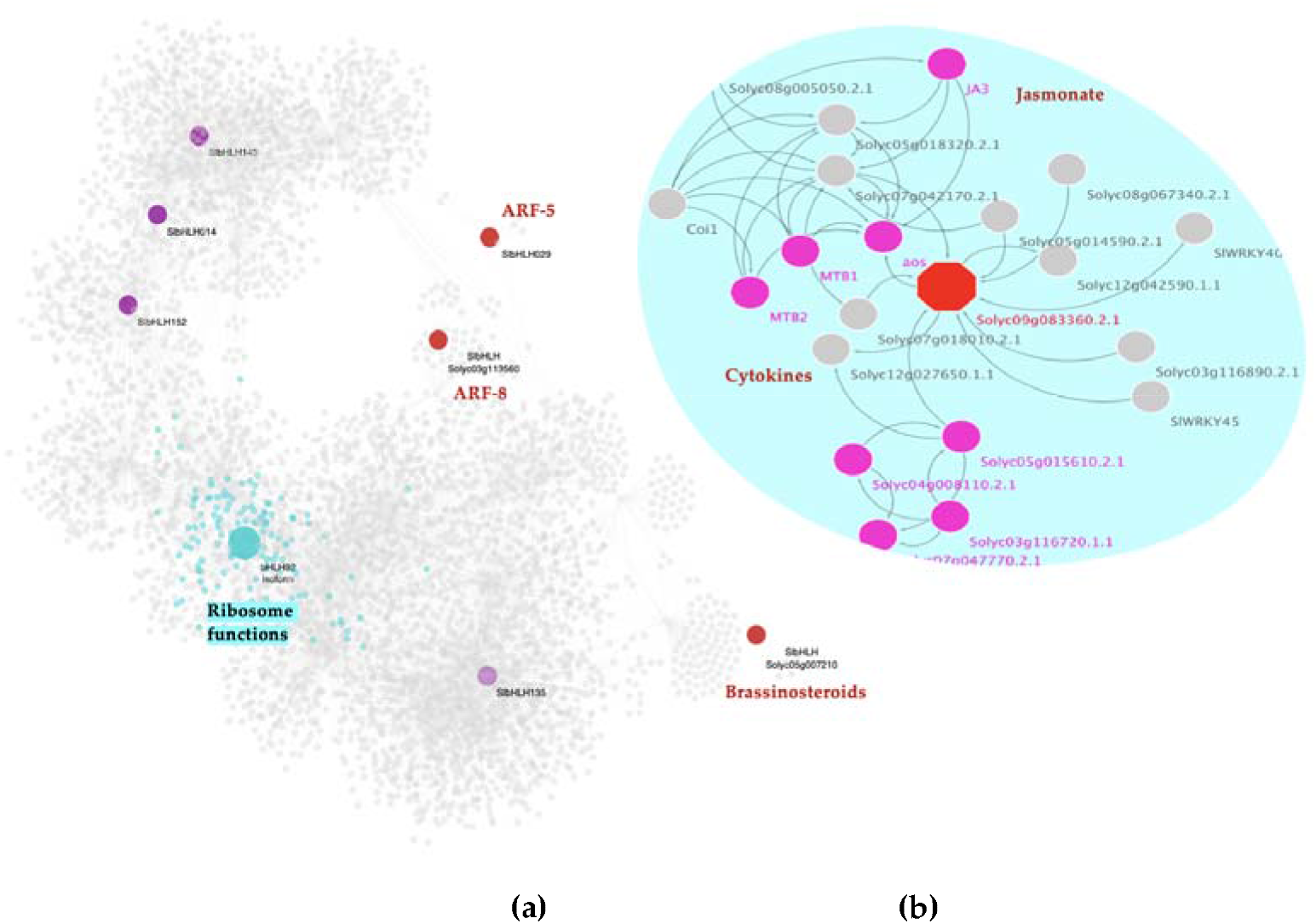
Microproteins predictive role in PSTVd severe infection: **(a)** Subnetwork depicting the presence of eight BHLH-TFs predicted as potencial microproteins. Nodes in red represent those that are related to hormone-events at a protein-protein interaction level. Node in cyan represents the BHLH-TF encoding the bhlh92 isoform identified as a MTR and its corresponding regulon (small nodes in the same color). Functional enrichment of its regulon is highlighted in cyan; **(b)** Protein-protein interaction network of the bhlh92 isoform *(Solyc09g083360.2.1)* the hexagon node represents this microprotein, while the fucsia nodes indicate the connected interactors related to jasmonate signaling (MYC2/JA3 BHLH transcription factor), while the others are linked to cytokines signaling events.

To deep insight into how these four hormones pathway-related miPs interact in each condition, we performed a mutual information algorithm analysis (26). These results suggest that bHLH miPs interactions in tomato hosts are distinctive among the three experimental conditions, **Figure S2**. Moreover, this analysis shows that these four biomolecules are present as regulators in the PSTVd-severe condition.

Strikingly, our MRA reveals that one of those miPs, bhlh92 isoform *(Solyc09g083360.2)* (NES= 2.2725 *p-value*= 0.02), is classified as a specific MTR for the S23 infected condition, At a genetic level, this BHLH-lF is a MTR that is positively regulating downstream 122 genes involved in ribonucleoprotein complex biogenesis and translation, among other ribosome-related functions, **Figure 5a**. While at a postranslational level, protein-protein interaction network reveals that bhlh92 isoform is a 233 aa protein with a predicted function related with aos a hydroperoxide dehydratase protein. Aos is a cytochrome P450 of the CYP74A subfamily member involved in the biosynthesis of jasmonic acid from lipoxygenase-derived hydroperoxides of free fatty acids. In a second interaction, it is linked to MYC2 also known as JA3 protein encoded by a BHLH TF, which we found as an MTR for the mild condition, **Figure 5b**. MYC2/JA3 is a transcriptional activator that binds to the G-box motif (5’-AACGTG-3’) found in the promoter of the jasmonate-induced gene LAPA1. It acts as a negative regulator of blue light-mediated photomorphogenesis and positively regulates root growth. This gene promotes growth in response to the phytohormones abscisic acid (ABA) and jasmonate (JA), and binds to the G-box motif (5’-CACGTG-3’) of the RBCS-3A gene promoter. This TF acts downstream of the jasmonate (JA) receptor to orchestrate JA-mediated activation of plant responses. It has been related to positive regulation of both wound-responsive and pathogen-responsive genes through MYC2 type genes.

In summary, the specificity of the presence of these biomolecules in the mutual interaction performed using the mild and severe infected samples, highlights that the differential expression and interaction of those miPs is very significant and could be relevant to the pathogenesis since its early development or mild symptoms.

### 2.8 BHLH, MYB, and ERF TF-families are MTRs at a tissue-specific level

In order to determine the tissue-specific regulation of the MTRs identified in the global integrated analysis, we analyze separately co-expression networks of leaf and root tissue samples. When performing the deconvolution of the network built with the integrated data (the global analysis), 69,022 interactions were obtained. While using only the expression data in the leaf transcriptomic dataset (specific tissue), a network of 29,316 interactions was obtained. The loss of interactions using tissue-specific samples compared to the global strategy is very noticeable.

Notably, accordingly with the global integrated analysis, in the leaves transcriptomic dataset, we identified a similar behavior of gene transcriptional regulation. We found that 15 out of the 21 MTRs identified in the C. vs S23 comparison of the global analysis are conserved in genetic regulation in leaf tissue. Moreover, MYB (11), and bHLH TFs (7) are the most representative families in the C. vs S23 comparison, in those specific samples. Interestingly, in this tissue-specific analysis the top MTR is a member of the ERF family the pathogenesis-related genes transcriptional activator, PTI6 *(Solyc06g082590.1)* which regulon is enriched in the KEGG:00073-Cutin, suberine, and wax biosynthesis synaptonemal complex organization.

When we deep insight into the tissue-specific regulation, we noticed that all the MTRs identified for the comparison between the PSTVd variants are also significant for the healthy-S23 comparison. This highlights the gene expression differences between the leaf cells infected within the two different PSTVd strains. Moreover, it is evidence of differences in their pathogenesis mechanisms. The top MTRs positions change according to the comparison, which in turn represents which regulons are most differentially expressed between the strains. For instance, the NAC TF family is highly relevant in the regulation by the severe variant in leaf tissue (7 members as MTRs); when comparing with the global analysis we find fewer members of this family acting as MTRs (3 members), **Figure 6a**.

**Figure 6.**
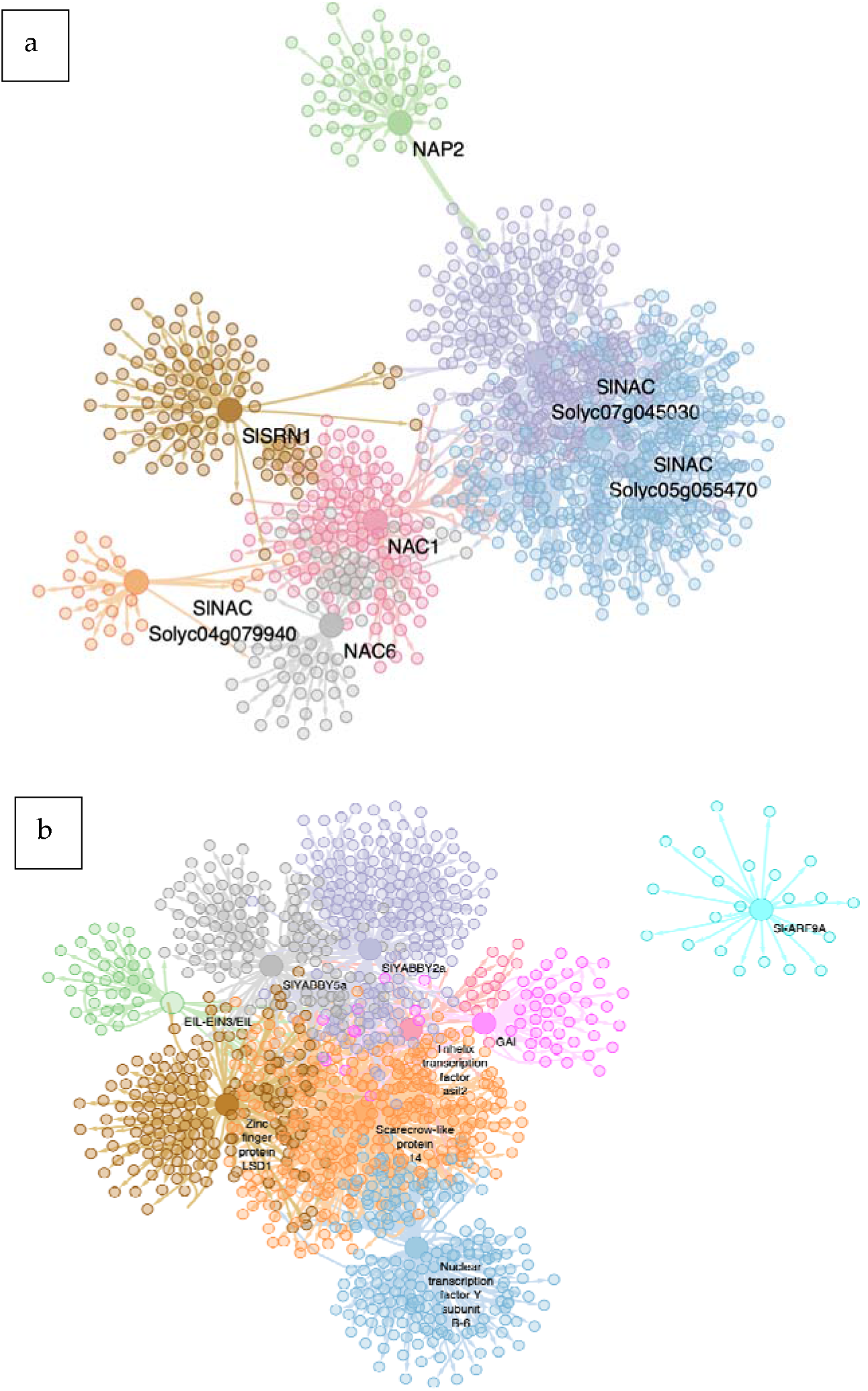
lnteractomes of the C vs S23 leaf tissue specific analysis. Each node represents a gene, the edges between nodes represent regulatory interactions between genes. The big highlighted nodes with a specific color represent MYB-MTRs, while nodes of the same color surrounding these represent target genes regulated by an MTR. (a) NAC family MTRs in tissue specific (leaf) Gene Regulatory Network, subnetworks where all the FTs of the NAC family that resulted as MTR for the C vs S23 comparison and their regulons are shown: SlSRN1 (brown), NAP2 (green), SlNAC Solyc07g045030 (purple), SlNAC Solyc05g055470 (blue), NAC1 (pink), NAC6 (gray), SlNAC Solyc04g079940 (orange). (b) Top MTRs in tissue specific (leaf) Gene Regulatory Network, subnetworks where top 10 highest NES MTRs for the C vs S23 comparison are shown: Nuclear Transcription Factor Y subunir-6 (blue), Scarecrow-like protein 14 (orange), Zinc finger protein LSDl (brown), EIL-EIN3/EIL (green), SIYABBY5a (gray), SIYABBY2a (purple) Trihelix TF asil2 (cherry red), GAI (pink), SI-ARF9A (cyan).

The TCP family is represented in the first two places at the top of the MTRs list for the C. vs S23 in our global analysis. However, in specific leaf tissue, they drop to positions 24 and 28, which indicates their loss of relevance in tissue-specific regulation. Additionally, other TFs families as MTRs appeared for the severe condition in the leaf-specific tissue analysis. These are YABBY TFs that are related to the regulation of the initial steps of embryonic shoot apical meristem (SAM) development, and in the abaxial cell fate determination during embryogenesis and organogenesis, respectively *(Solyc07g008180.2, Solyc06g073920.2)* [51–52], GRAS *(Sol ycl l g011260.1, Sol yc05g053090.1)* transcriptional regulators that act as repressors of the gibberellin (GA) signaling pathway. They probably act by participating in large multiprotein complexes that repress transcription of GA-inducible genes, EIL (Solyc01g009170.2)binds a primary ethylene response element present in the ETHYL EN E-RESPO NSE-FACTOR} promoter with consequence to activate the transcription of this gene [53], LSD a positive regulator of reactive oxygen-induced cell death *(Solyc08g077060.2)*, NF-YB a Histone-fold superfamily member *(Solyc06g069310.2)*, Trihelix *(Solyc01g096470.2)*, and ARF an auxin-activated signaling pathway *(Solyc08g082630.2)*, **Figure 6b**.

In contrast, when we performed the same analysis using the root transcriptomic data none MTR was identified. This could be due to the more evident gene differential expression among samples in the leaves microarray than in the roots transcriptomic data, where samples expression remain highly similar, **Figure S4**. In this sense, when discarding the root samples, the amount of the MTRs for all comparisons is larger in leaf-specific tissue than in the global analysis. In the C. vs S23 123/59 MTRs (tissue-specific/global, respectively), C. vs M 25/7, and M vs S23 61/21 (threshold *p=0.05)*. Overall, our results from tissue-specific networks analysis reveal the conservancy of the pivotal role of the bHLH, MYB, and ERF families at a tissue-specific level in the tomato response to the PSTVd infection.

## 3. Discussion

Viroid pathogenesis relies upon highly complex molecular and biological processes orchestrated by both the host and the viroid genomes. This interplay of events results in asymptomatic, mild, or severe symptoms. Taking advantage of the recent time-course transcriptome analysis of the PSTVd-tomato interaction we delved into the transcriptional regulation of genes coordinated by MTRs. It is well known that transcriptional reprogramming is governed by MTRs. In this context, it is crucial to delineate the pivotal role played by those regulators in coordinating specific regulatory networks. Moreover, the role of MTRs in the viroid pathogenesis mechanism is still unknown.

Using an *omics* approach that integrates PSTVd-infected tomato transcriptomic datasets, and the *corto* algorithm, we found the most important TFs regulating gene expression of the given phenotype of a plant-pathogen caused disease. By comparing the healthy condition (C) with the abnormal phenotypes (S23 and M) we determined which regulators could be responsible for the physiological changes underlying viroid infection. Identifying the target genes of the inferred MTRs helps us to functionally characterize them for a better comprehension of how these gene expression alterations are related to the different representative symptoms of the condition of interest. Furthermore, comparing two different symptom phenotypes can provide us with an insight into the differences between the two phenotypes and the different mechanisms underlying the pathogenesis displayed by the two strains belonging to the same pathogen species, in this case, PSTVd.

In a global vision, the mechanism of viroid pathogenesis is highly complex, it involves the concerted interplay among host factors, as an outcome of a mature-viroid/host factor interaction, activating protein kinases, and pathogenesis reponse proteins (PRs), as well as the recruitmeint of enzymes to accomplished their replication, and movement. Other mechanism relies on the activation of hormone-related responses, where microproteins and MTRs may be key molecules in the impairment of protein functions, changes in miRNAs and siRNAs pathways. Additionally, viroid intermediates from replication can be loaded into the RNA induced complex (RISC) to generate vd-siRNAs that are capable to guide mRNA degradation. Overall, those events are coupled together to modularte the host gene expression, and are dependent upon the host/ viroid strain combination, as well as on the participation of enviromental factors. As an outcome of those highly specific and regulated events, a tolerant or a susceptible phenotype will be displayed in infected plants, **Figure 7**. At the plant cell level, the key molecules are regulating genetic, epigenetic, and post-translational events in the nucleus, cytoplasm and ribosomes. For instance, the mature viroid is interacting with RNA polymerase II and other proteins for its replication, microproteins are forming multi protein complexes to negatively regulate transcription, MTRs are positively or negativily regulation downstream genes, and vd-siRNAs are pathogenic effectors that act guiding the degration of mRNA in postranslational ways, all these interactions can be studied using networks approaches.

**Figure 7.**
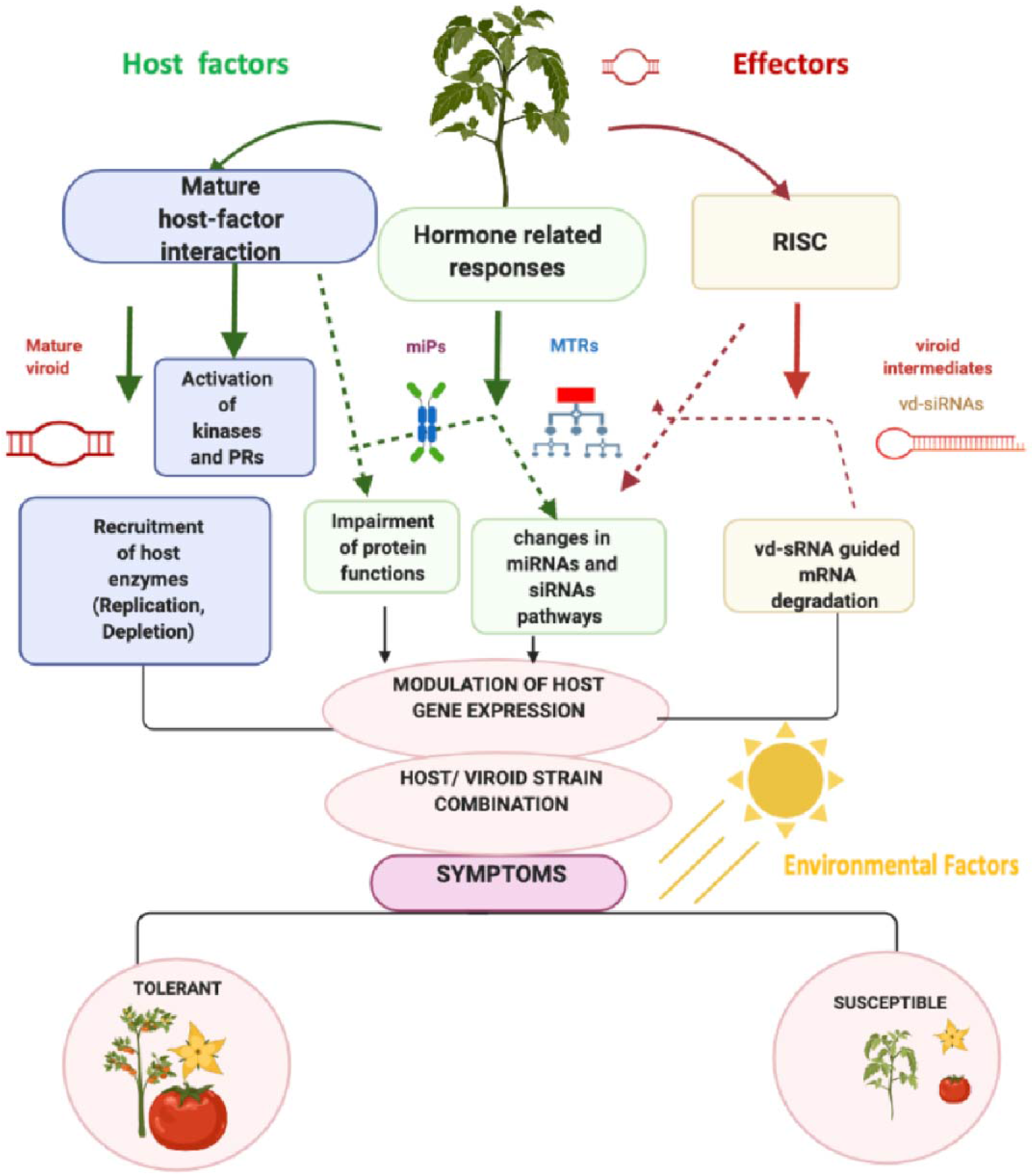
Proposed mechanism of viroid-host interplay, symptom development is an outcome of the complex plant-pathogen interaction as a result of alterations to cell processes related to viroid RN A-mediated genetic regulation.

In this study, we implemented a computational strategy to identify and classify the MTRs, the functional role of their regulons, as well as potential functional microproteins under the described experimental conditions. An outstanding example of how this computational strategy differentiates at the viroid strain level is by identifying those key molecules that are conserved when comparing the PSTVd-S23 strain infected samples vs. the mock-inoculated control plants; as well as the comparison of the M strain infected vs. the S23 infected plants. For instance, is clearly apparent in the MYB-TFs family, where *kua1 isoform x1(Solyc08g078340.2*) MTR is conserved for these two comparisons. This suggests that the differential expression caused by the severe strain is very strong and that even with an infection with a strain of the same viroid species that generates moderate symptoms (M. which can be attributed to a differential expression of lesser magnitude), a significant difference is still observable. Moreover, this conservation of gene transcriptional regulation can be seen in the members of the ERF-TFs family. In this family, two conserved members act as MTRs when comparing C. vs. S23 and M. vs. S23 AP2/EREBP TF1 *(Solyc02g093130.1)*, and ERF_C_5 *(Solyc02g077370.1)*. Also, this result indicates that their differential expression is still appreciable when comparing samples of both the mild and severe PSTVd variants. It is also evident from the data analysis that there are five regulators with no differential expression (absent) when comparing the M. vs. S23 conditions ERF-1a *(Solyc05g051200.1), RAP2-12(Solyc12g049560.1)*, PTI6 *(Solyc06g082590.1)*, TSRF1 *(Solyc09g089930.1)*, and ERF_A_2 *(Solyc03g093610.1)*. One conserved MYB-MTR can be differentiated between the PSTVd variants. These results are in accordance with the previous PSTVd-root and leaf tissue-specific transcriptomic analysis, where a similar behavior of MYB-TFs regulating gene expression as a host responsive mechanism under infection by the two variants was previously reported. These results demonstrate the potential of the approach we developed for this plant-overall integrated study where additionally we identified the potential downstream target genes of each of the MTRs.

As has been previously reported, we found that the major TF families with differential expression in the S23 and M infection are bHLH and ERF-types which play important roles in plant defense [7,15]. In root tissue, 50% of the highest up-regulated genes are shared in infections from both PSTVd-strains. Among the shared genes are those coding for bHLH-family TF, which are transcriptional co-repressors of genes encoding glyoxylate reductase, NAD kinase, calcium-transporting ATPase, and chlorophyll a-b binding proteins, among other functions. Additionally, in our analysis, we included the ERF family. Its role was characterized due to its abundance in plants expressing severe symptoms. Moreover, changes in the expression of TFs belonging to this family were formerly reported for viroid infection [3,7,15]. Members of this TF family are crucial in the regulation of plant development and growth, fruit ripening, defense response, and metabolism. ERFs are modulators of hormone-related processes that involve ethylene, gibberellins, cytokinins, and abscisic acid biosynthesis in response to auxin, cytokinin AUX, CK, GB, and ABA [27–29,39,42–43].

We also identified those MTRs which are distinct or unique for each infection condition. Examples of this are also observable for the MYB-TFs family. We identify two members of this family classified as unique MTRs for the C. vs S23 comparison, MYBlRl *(Solyc04g005100.2)*, and MYB16 *(Solyc02g088190.2)*, and those were absent for the M. vs S23 comparison. Thus, this indicates that there is also a high differential expression of them in mild strain infection, resulting in similar levels to those of the severe strain infection. Notably, one MYB-TF was identified as a unique MTR when comparing M. vs S23, PHR1-LIKE 1*(Solyc05g055940.2)*, and this TF may function as a differential regulator of symptom development induced by the two strains.

A totally opposite case occurs in the regulation of tomato gene expression by the bHLH TF family during viroid infection. In this TF family, MTRs identified in the two strain comparisons are all different, highlighting that none of them is conserved. This could suggest that the two PSTVd variants undergo transcriptional reprogramming via the same bHLH-TFs as MTRs to establish the mechanism of pathogenesis (such as SlbHLH0ll *(Solyc01g111130.2)*, bHLH130-like *(Solyc12g100140.1)*, SlbHLH022 *(Solyc03g097820.1)*, GBOF-1*(Solyc06g072520.1)*, and bhlh92 isoform *(Solyc09g083360.2)*. In accordance with recent studies that identified transcription factors differentially expressed for mild and severe infection in two pepper cultivars, our results show that MYB-TFs, Homeodomain, WRKY, and Heat-Shock-related are specific for the severe condition [7].

Since it is impossible to describe an exhaustive list of all the TFs and their potential roles in one communication, we have performed an independent study that adopted the co-expression modularity method, [54]. The objective is to characterize co-expression modules of the major TF families, including members of the bHLH, that display indispensable roles in the plant immune response. For instance, the MYC2-TF is likely to be a hub gene for the development of symptoms (24dpi) displayed, particularly by the PSTVd-mild strain, [55].

In accordance with other studies that described the relevance of pathways involved in carbohydrate metabolism, we found a high enrichment of the regulons of bHLH, ERF, and unique MTRs that are predicted to be participating together in the negatively transcriptional reprogramming for the severe strain condition.

A recent study of gene expression in CBCV infected commercial hop highlighted the involvement of genes associated with the Brassinosteroid pathway, as evidenced by the profound protein-protein interactions (PPI) of hub genes such as BIM1 and BIM2 with the hop bHLH-TFs, [56]. In our study, we approach the potential functional role of members of this family in the tomato genome as miPs candidates. We identified four bHLH-TFs that are interacting with high relevance in the PSTVd-severe condition. To the best of our knowledge, this represents the first approach to study the role of miPs in viroid infection.

Additionally, it is noteworthy that other TFs identified in our study could be relevant and must be studied in more depth. Recently we have identified that members of a novel and poorly described TF family (PLATZ-TFs) are potential candidates as targets of regulation by multiple tomato planta macho viroid (TPMVd)-vdsiRNAs, [57]. Holistic approaches focusing on complex interactions within biological systems are now being employed to study viroid-host interactions at different regulatory levels, including epigenetic, genomic, and small RNA interference [58]. Further studies that perform multi-omics analysis at the level of the genome host (key molecules) and pathogen interactors (effectors such as vd-siRNAs) will help delineate future strategies related to the use of these biomolecules which could serve as a basis for further analysis in the journey of the development of improved management for this agronomically important diseases.

## 4. Materials and Methods

### 4.1 Designed pipeline for the omics approach

For this study, we implemented a pioneering method that integrates transcriptomic data for co-expression and regulatory network analysis to give deep insight into the molecular mechanisms underlying *Solanum lycopersicum* response to PSTVd infection, **Figure S3**. First, we performed a transcriptomic integrative data analysis, followed by a step of normalization and data processing in order to have a unique expression matrix (Steps 1-2). Then, we used the PlantTFDB (http://planttfdb.gao-lab.org/) to obtain TF lists, [31]. To perform the interactomics analysis, we used those TF-lists, and the unique expression matrix as inputs. We carried out network deconvolution analysis to have GRN as outputs using the corto algorithm(Steps 3-4) [59–60]. Subsequently, functional enrichment analysis of the regulons under study was performed (Step 5). Finally, the miPs assignment and the PPI analyses were carried out (Step 6).

### 4. Microarray Data Integration and TF Datasets for the GRN

We obtained two datasets describing transcriptome-wide effects of PSTVd-tomato infection. Microarray expression datasets of root (GSE111736) and leaf (GSE106912) tomato transcriptome in mild and severe PSTVd infection were obtained from the NCBI GEO database (https://www.ncbi.nlm.nih.gov/gds). Those studies consisted of experiments employing a time-course in three stages using Control (C), PSTVd-mild (M), and PSTVd-severe (S23) samples. The time-course is as follows: early symptoms (17 dpi) samples, complete symptoms (24 dpi), and recovery (49 dpi). In total, 26 root and 27 leaf samples were separately analyzed, [7, 15]. In our study, data integration of both transcriptome datasets was performed to obtain a unique expression matrix. Data integration consisted in merging datasets in order to have only the variant infection treatments of both tissues. A mean value was calculated for each repeated gene on the dataset, then the matrix was normalized using the MRA function of the affy package, [59]. Our integrated expression matrix is composed of a total of 53 comparable samples into three different conditions: C: 18 samples 9 from each tissue; PSTVd-M: 18 samples, 9 from each tissue; and PSTVd-S23: 17 samples 9 and 8 from each tissue.

### 4.3 Identification of biological communities in the Gene Regulatory Network

To approach the study of the biological significance of the genes in the regulatory networks, they were grouped, according to their role in biological communities. This analysis was estimated using the Louvain method. The Louvain method is a greedy optimization method that optimizes the modularity of a partition of the network [48]. These metrics were implemented using the NetworkX package in Python scripts, [61]. Then, functional enrichment analyses were assessed to each estimated community using the over-representation analysis of the functional assignment terms, including GO: terms and KEGG pathways. The estimated communities that do not have functional relevance were discarded. The g: Profiler R package was employed for data analysis, [62].

### 4.4 Identification of bHLH-miPs and their functional assignment

For the miPs classification, we employed the miPFinder program (https://github.com/DaStraub/miPFinder) using the bHLH TF list from the PlantTFDB to determine which members are potential candidates for encoding functional microproteins. Then, protein-protein interaction analyses were performed to each estimated bHLH-miP using the String database (https://string-db.org/). The miPs that do not have functional relevance linked to hormone pathways were discarded. For each miP-ppi network visualization, the Cytoscape environment was used, [63].

### 4.5 Deconvolution of the Gene Regulatory Network

For the inference of the coexpression-based networks the corto algorithm with default parameters, freely available on the CRAN repository of R packages, was used. The corto package is a co-expression-based tool that infers GRNs using a TFs list with their targets, and an expression matrix data set, [60]. In brief, corto uses a combination of Spearman correlation and Data Processing Inequality (DPI), adding bootstrapping to evaluate the significant edges, removing indirect interactions. As an input, we used the integrated expression matrix and the TF list described in the <u>4.1 Microarray Data Integration and TF Datasets for the GRN</u> section. For this analysis, a *p-value*= 1 × 10-8 was used as a cut-off, and 100 bootstraps were carried out. Subsequently, a single network for the integrated expression matrix was obtained, generating, as a result, an output file in which an inferred enriched GRN is contained.

### 4.6 Identification of the Master Transcriptional Regulators (MTRs) and their functional relevance in the three experimental conditions

Master Regulator Analyses were performed by comparing infected and mock samples with the corto algorithm, using default parameters and the coexpression network derived from the integrated expression dataset. Using this R package we inferred the master transcriptional regulators in the transition between two given conditions.The MRA function implicit in the package calculates the enrichment of each TF-centered network in a user-selected signature (a list specifying which samples correspond to each condition), provided as two gene expression matrices (eg, diseased tissue vs. control). For this study, we performed an MRA for each comparison among conditions; S23 vs. C.; M. vs. C; S23 vs. M. In order to determine the master transcriptional regulators, we compared among those three different conditions using a *p-value* of 0.05 as a cut-off. As an output, we obtain a list of master regulators ordered by their Normalized Enrichment Score (NES), which is a measurement of how different is the expression of a given regulon when comparing two conditions. A combined value for the coexpression network is generated by weighing every gene likelihood in the network, providing a final NES, which is positive if the regulon is upregulated by the infection, and negative if it is downregulated. Finally, functional enrichment analyses were assessed to each regulon estimated using the over-representation analysis of the functional assignment terms, including GO: terms and the KEGG pathways. The g: Profiler R package was employed for data analysis, [62]. For each subnetwork visualization, the Cytoscape environment was used, [63].

## 5. Conclusions

Gene regulatory networks of Mild, and S23-severe PSTVd strain samples from transcriptomics analysis reveal that specific bHLH, MYB, and ERF master transcriptional factors are regulating genes among experimental treatments. By performing network analysis we determine those MTRs and their targets that participate in molecular mechanisms underlying distinct and shared biological processes. Notably, miPs characterization in networks among treatments pointed out that bHLH TFs are potential miPs involved in the post-translational regulation which suggests being distinct. For instance, the severe PSTVd strain is related to auxin-responsive factors as well as brassinosteroid and jasmonate hormones. Our study represents a pioneer attempt to identify and classify PSTVd-responsive tomato master transcriptional regulators and provides valuable information about their involvement in the development of viroid pathogenesis in this host. Altogether, our results lay a foundation for further research on the PSTVd and host genome interaction, providing evidence for identifying potential key genes that influence symptoms during development in tomato plants.

## Supplementary Materials

**Figure S1.**
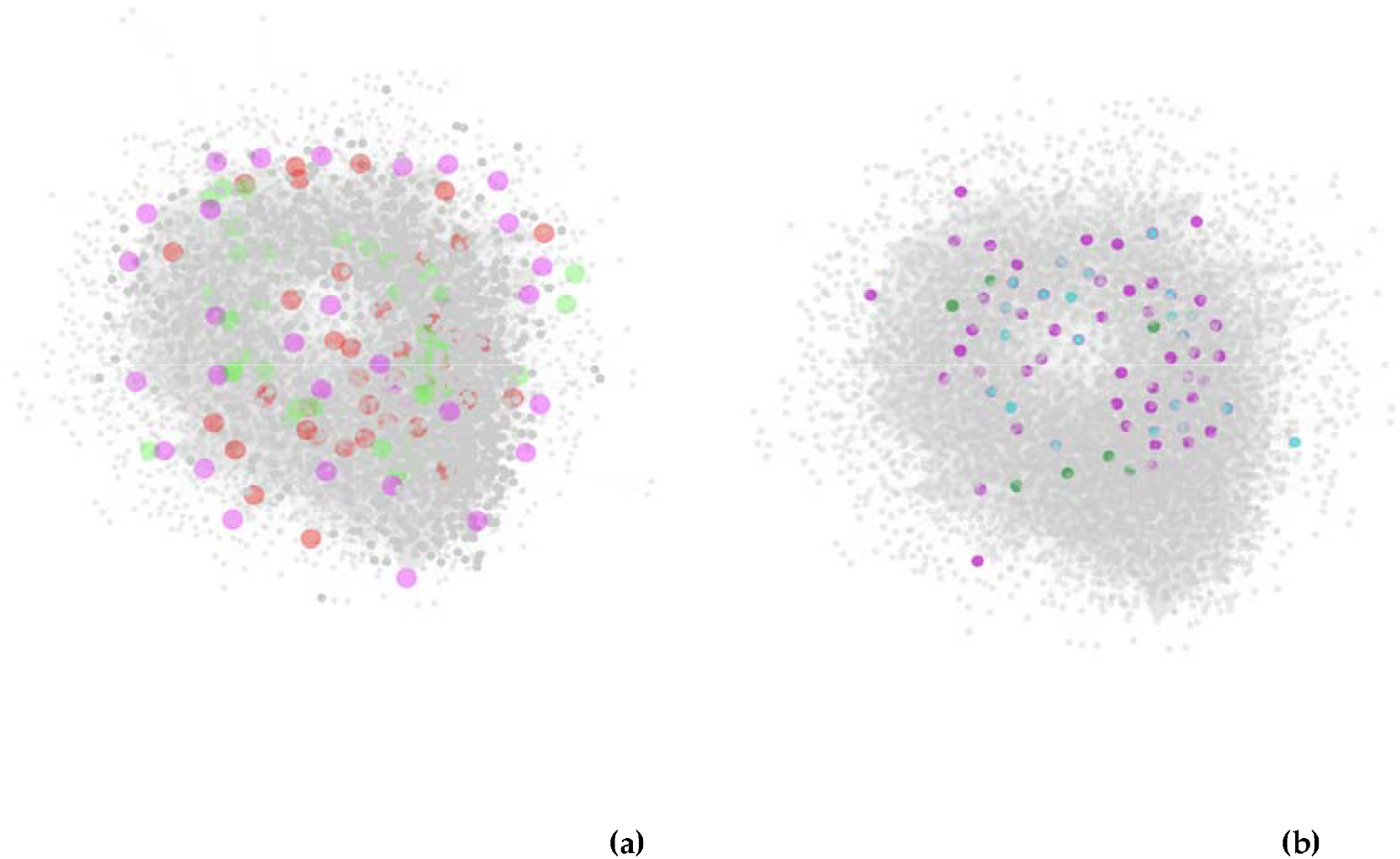
Gene Regulatory Global Network. (a) each node represents a gene; the edges among nodes depict the regulatory action of one node on another(s). The most prominent nodes highlighted in different colors indicate all the TFs of the families of interest in this study that are present in the global network; bHLH-TF (pink), MYB-TF (green), and ERF-TF (red),(b)Interactome of the 87 MTRs of the TFs of of interest; C vs S23 (purple), C vs M (green), and M vs S23 (blue).

**Figure S2.**
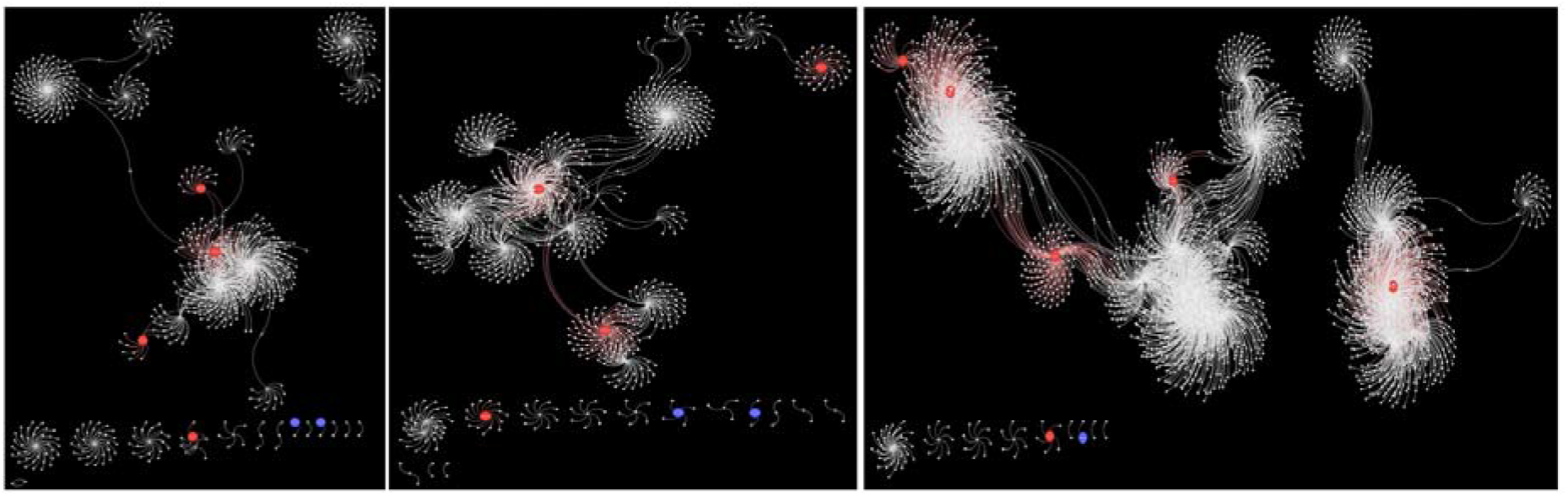
Mutual interaction Networks. miP-molecule interactions among the different conditions from left to right a) mock-inoculated samples, b) PSTVd-mild strain, c) PSTVd-S23. Each node represents a gene, the edges between nodes depicts putative interactions among them(regulation). Nodes highlighted in red color represent the hub-miPs, while the edges of the same color represent their interactions with other molecules. Nodes in blue color represent miPs.

**Figure S3.**
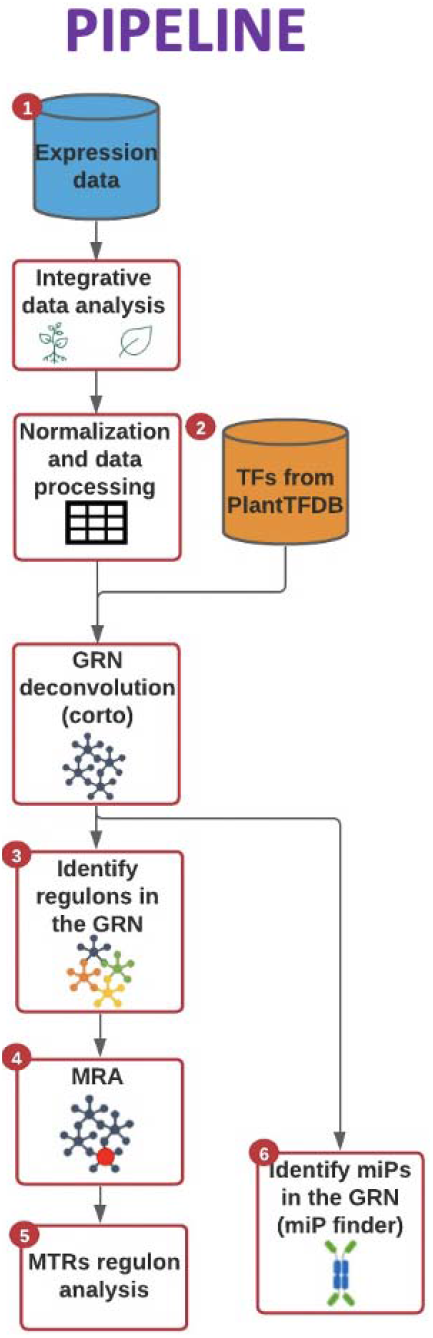
Pipeline for the omics approach. 1) First, microarray publicly available expression data of both control and infected leaves and roots were processed to obtain a unique expression matrix. 2) The expression matrix and a list of tomato genome transcription factors from PlantTFD were used as input to infer a network of transcriptional regulation employing Corto alghoritm. 3)Then, a regulon network was obtained by associating the expression level of the targets of all the transcription factors. 4) Subsequently, the MRA algorithm was used to calculate the MTRs in the network. 5) Once the MTRs are infered we proceeded to explore the functional role of them and their regulons. 6) Additionally, the bHLH-coding genes for miPs were assigned in the tomato genome. Among them, we identify those that are interacting in the GRN, and those that could be potential MTRs.

**Figure S4.**
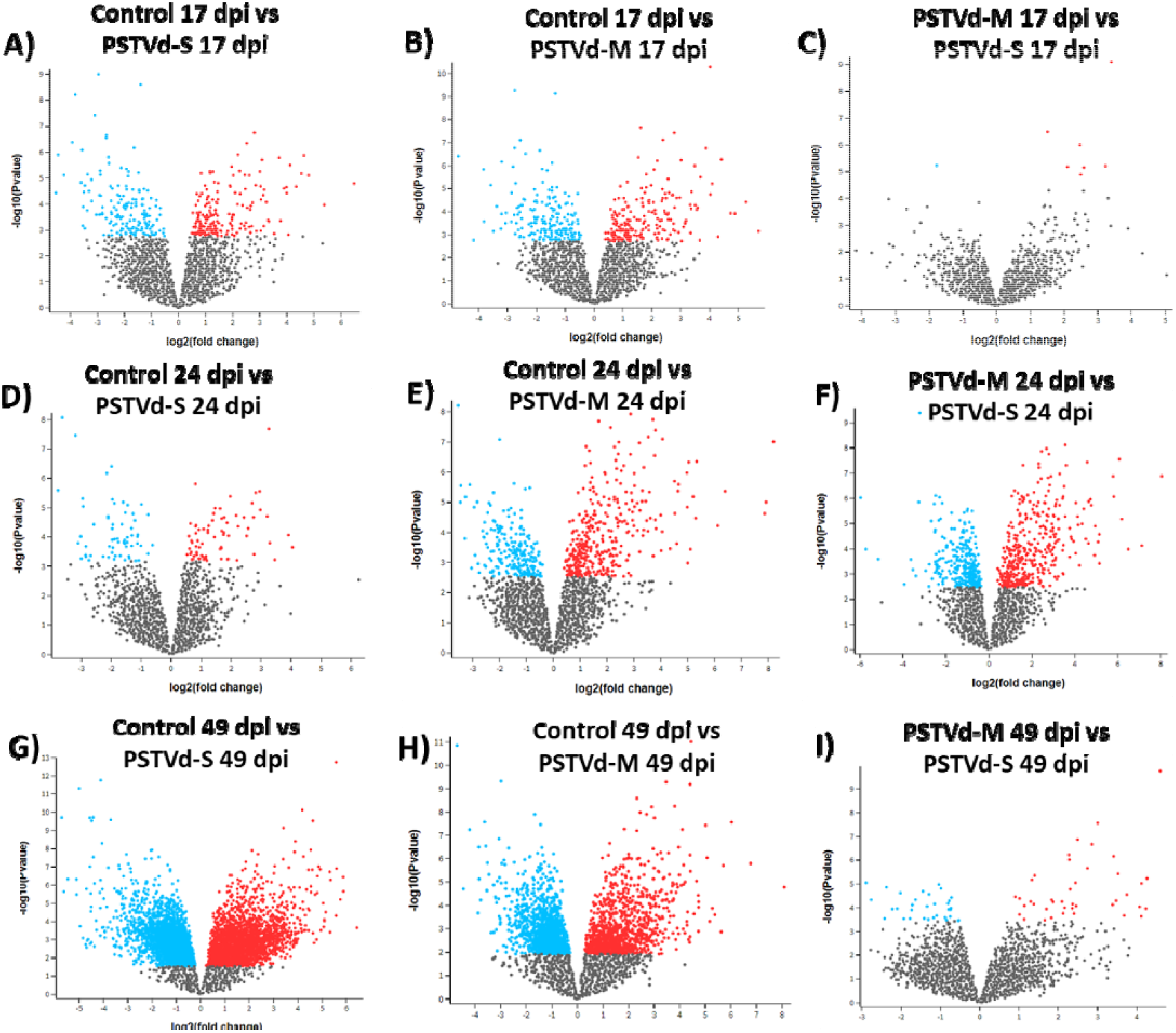
Comparative analysis of gene expression among treatments in tomato roots. Volcano plots depicting the differential expression among control, severe PSTVd-S and mild PSTVd-M strains in the three phases of infection. Early (17 dpi), complete symptoms (24 dpi), and recovery (49 dpi). Each dot represents one gene and red color dots denote significant differentially expressed genes that achieved adjusted *p-value* <0.05.

## Author Contributions

conceptualization, K.A.P, R.W.H., and M.H.R; Methodology, K.A.P., and M.H.R; Bioinformatics analysis, K.A.P, O.M.Z, GE.HO, P.A, and M.A.J.L; Resources, K.A.P, M.H.R; Writing the original draft, K.A.P, and O.M.Z.; Writing, reviewing, and editing, K.A.P., P.A., M.H.R, and R.W.H; Data visualization, K.A.P, G.E.H.O, and O.M.Z; Supervision, K.A.P., M.H.R., and R.W.H.; Funding acquisition, K.A.P, M.H.R., and R.W.H. All authors have read and agreed to the published version of the manuscript.

## Data Availability Statement

### Datasets

Microarray from roots samples: https://www.ncbi.nlm.nih.gov/geo/query/acc.cgi?acc=GSEl11736

Microarray from leaf samples: https://www.ncbi.nlm.nih.gov/geo/query/acc.cgi?acc=GSE106912

Integrated expression matrix from both tissues: https://github.com/MarcoJL9/Network-analysis-of-root-and-leaf-transcriptome-integration

Code: https://github.com/MarcoJL9/Network-analysis-of-root-and-leaf-transcriptome-integration

## Acknowledgments

We would like to thank Dr. Aneta Więsyk for her support in providing the datasets used in the current paper. For technical support we thank Felipe de Jesus Rodriguez Gaxiola. For critical suggestions, we thank Dr. Hugo Antonio Tovar Romero.

## Appendix A. List of the gene communities found in the BHLH, ERF, and MYB interactomes

https://docs.google.com/spreadsheets/d/1kpS7nP7MtCda9TIFZX7xboo9lDjBcPz7qABIOQHFROA/edit#gid=0

## Appendix B. Functional enrichment analysis of the identified regulons of theMTRs

## References

1. Diener, T. O. Potato Spindle Tuber “Virus.” Virology 1971, 45 (2), 411–428. https://doi.org/10.1016/0042-6822(71)90342-4.

2. Katsarou, K.; Adkar-Purushothama, C. R.; Tassios, E.; Samiotaki, M.; Andronis, C.; Lisón, P.; Nikolaou, C.; Perreault, J.-P.; Kalantidis, K. Revisiting the Non-Coding Nature of Pospiviroids. Cells 2022, 11 (2), 265. https://doi.org/10.3390/cells11020265.

3. Aviña-Padilla, K.; Rivera-Bustamante, R.; Kovalskaya, N.; Hammond, R. Pospiviroid Infection of Tomato Regulates the Expression of Genes Involved in Flower and Fruit Development. Viruses 2018, 10 (10), 516. https://doi.org/10.3390/v10100516.

4. Wang, Y.; Shibuya, M.; Taneda, A.; Kurauchi, T.; Senda, M.; Owens, R. A.; Sano, T. Accumulation of Potato Spindle Tuber Viroid-Specific Small RNAs Is Accompanied by Specific Changes in Gene Expression in Two Tomato Cultivars. Virology 2011, 413 (1), 72–83. https://doi.org/10.1016/j.virol.2011.01.021.

5. Owens, R. A.; Tech, K. B.; Shao, J. Y.; Sano, T.; Baker, C. J. Global Analysis of Tomato Gene Expression during Potato Spindle Tuber Viroid Infection Reveals a Complex Array of Changes Affecting Hormone Signaling. Mol Plant Microbe Interact 2012, 25 (4), 582–598. https://doi.org/10.1094/mpmi-09-11-0258.

6. Itaya, A.; Matsuda, Y.; Gonzales, R. A.; Nelson, R. S.; Ding, B. Potato Spindle Tuber Viroid Strains of Different Pathogenicity Induces and Suppresses Expression of Common and Unique Genes in Infected Tomato. Mol Plant Microbe Interact 2002, 15 (10), 990–999. https://doi.org/10.1094/mpmi.2002.15.10.990.

7. Więsyk, A.; Iwanicka-Nowicka, R.; Fogtman, A.; Zagórski-Ostoja, W.; Góra-Sochacka, A. Time-Course Microarray Analysis Reveals Differences between Transcriptional Changes in Tomato Leaves Triggered by Mild and Severe Variants of Potato Spindle Tuber Viroid. Viruses 2018, 10 (5), 257. https://doi.org/10.3390/v10050257.

8. Hadjieva, N.; Apostolova, E.; Baev, V.; Yahubyan, G.; Gozmanova, M. Transcriptome Analysis Reveals Dynamic Cultivar-Dependent Patterns of Gene Expression in Potato Spindle Tuber Viroid-Infected Pepper. Plants 2021, 10 (12), 2687. https://doi.org/10.3390/plants10122687.

9. Wang, Y.; Wu, J.; Qiu, Y.; Atta, S.; Zhou, C.; Cao, M. Global Transcriptomic Analysis Reveals Insights into the Response of “Etrog” Citron (Citrus Medica L.) to Citrus Exocortis Viroid Infection. Viruses 2019, 11 (5), 453. https://doi.org/10.3390/vl1050453.

10. Tessitori, M.; Maria, G.; Capasso, C.; Catara, G.; Rizza, S.; De Luca, V.; Catara, A.; Capasso, A.; Carginale, V. Differential Display Analysis of Gene Expression in Etrog Citron Leaves Infected by Citrus Viroid III. Biochim Biophys Acta 2007, 1769 (4), 228–235. https://doi.org/10.1016/j.bbaexp.2007.03.004.

11. Herranz, M. C.; Niehl, A.; Rosales, M.; Fiore, N.; Zamorano, A.; Granell, A.; Pallas, V. A Remarkable Synergistic Effect at the Transcriptomic Level in Peach Fruits Doubly Infected by Prunus Necrotic Ringspot Virus and Peach Latent Mosaic Viroid. Virol J 2013, 10 (1). https://doi.org/10.1186/1743-422x-10-164.

12. Kappagantu, M.; Bullock, J. M.; Nelson, M. E.; Eastwell, K. C. Hop Stunt Viroid: Effect on Host (Humulus Lupulus) Transcriptome and Its Interactions with Hop Powdery Mildew (Podospheara Macularis). Mol Plant Microbe Interact 2017, 30 (10), 842–851. https://doi.org/10.1094/mpmi-03-17-0071-r.

13. Xia, C.; Li, S.; Hou, W.; Fan, Z.; Xiao, H.; Lu, M.; Sano, T.; Zhang, Z. Global Transcriptomic Changes Induced by Infection of Cucumber (Cucumis Sativus L.) with Mild and Severe Variants of Hop Stunt Viroid. Front Microbiol 2017, 8. https://doi.org/10.3389/fmicb.2017.02427.

14. Mishra, A.; Kumar, A.; Mishra, D.; Nath, V.; Jakše, J.; Kocábek, T.; Killi, U.; Morina, F.; Matoušek, J. Genome-Wide Transcriptomic Analysis Reveals Insights into the Response to Citrus Bark Cracking Viroid (CBCVd) in Hop (Humulus Lupulus L.). Viruses 2018, 10 (10), 570. https://doi.org/10.3390/v10100570.

15. Góra-Sochacka, A.; Więsyk, A.; Fogtman, A.; Lirski, M.; Zagórski-Ostoja, W. Root Transcriptomic Analysis Reveals Global Changes Induced by Systemic Infection of Solanum Lycopersicum with Mild and Severe Variants of Potato Spindle Tuber Viroid. Viruses 2019, 11 (11), 992. https://doi.org/10.3390/vll110992.

16. Guzzi, P. H.; Mercatelli, D.; Ceraolo, C.; Giorgi, F. M. Master Regulator Analysis of the SARS-CoV-2/Human Interactome. J Clin Med 2020, 9(4), 982. https://doi.org/10.3390/jcm9040982

17. Ghazalpour, A.; Doss, S.; Zhang, B.; Wang, S.; Plaisier, C.; Castellanos, R.; Brozell, A.; Schadt, E. E., Drake; T. A.; Lusis, A. J.; Horvath, S. Integrating genetic and network analysis to characterize genes related to mouse weight. PLoS Genet 2006, 2(8), e130. https://doi.org/10.1371/journal.pgen.0020130

18. Bonin-Andresen, M.; Smiljanovic, B.; Stuhlmüller, B.; Sörensen, T.; Grützkau, A.; Häupl, T. Bedeutung von Big Data für die molekulare Diagnostik [Relevance of big data for molecular diagnostics]. Z Rheumatol 2018, 77(3), 195–202. https://doi.org/10.1007/s00393-018-0436-3

19. Souza, G.; Salvador, E. A.; de Oliveira, F. R.; Cotta Malaquias, L. C.; Abrahão, J. S.; Leomil Coelho, L. F. An in silico integrative protocol for identifying key genes and pathways useful to understand emerging virus disease pathogenesis. Virus Res 2020, 284, 197986. https://doi.org/10.1016/j.virusres.2020.197986

20. van Someren, E. P.; Wessels, L. F. A.; Backer, E.; Reinders, M. J. T. Genetic Network Modeling. Pharmacogenomics 2002, 3 (4), 507–525. https://doi.org/10.1517/14622416.3.4.507.

21. Kuijjer, M. L.; Tung, M. G.; Yuan, G.; Quackenbush, J.; Glass, K. Estimating Sample-Specific Regulatory Networks. iScience 2019, 14, 226–240. https://doi.org/10.1016/j.isci.2019.03.021

22. Spirtes, P.; Glymour, C.; Scheines, R.; Kauffman, S.; Aimale, V.; Wimberly, F. Constructing Bayesian Network Models of Gene Expression Networks from Microarray Data. Carnegie Mellon University. Journal contribution 2000. https://doi.org/10.1184/Rl/6491291.vl

23. Ovens, K.; Eames, B. F.; McQuillan, I. The impact of sample size and tissue type on the reproducibility of gene co-expression networks. Proceedings of the 11th ACM International Conference on Bioinformatics, Computational Biology and Health Informatics 2020. New York, NY, USA, Article 12, 1–10. DOI:https://doi.org/10.1145/3388440.3412481

24. Hernández-Lemus, E.; Tovar, H. Networks of Transcription Factors. Genome Plasticity in Health and Disease 2020, 137–155. https://doi.org/10.1016/b978-0-12-817819-5.00009-7.

25. Kimura, S.; Sinha, N. Tomato (Solanum Lycopersicum): A Model Fruit-Bearing Crop. CSH Protoc 2008, 2008 (12), pdb.emo105–pdb.emo105. https://doi.org/10.1101/pdb.emo105.

26. Maureira, F.; Rajagopalan, K.; Stöckle, C. O. Evaluating Tomato Production in Open-Field and High-Tech Greenhouse Systems. J Clean Prod 2022, 130459. https://doi.org/10.1016/j.jclepro.2022.130459.

27. FAO. FAOSTAT Food and Agricultural Organization Statistics http://faostat.fao.org/site/567/DesktopDefault.aspx?PageID=567#ancor (accessed Jun 16, 2021).

28. Ling, K.; Zhang, W. First Report of a Natural Infection by Mexican Papita Viroid and Tomato Chlorotic Dwarf Viroid on Greenhouse Tomatoes in Mexico. Plant Dis 2009, 93 (11), 1216–1216. https://doi.org/10.1094/pdis-93-11-1216a.

29. Ling, K.-S.; Verhoeven, J. Th. J.; Singh, R. P.; Brown, J. K. First Report of Tomato Chlorotic Dwarf Viroid in Greenhouse Tomatoes in Arizona. Plant Dis 2009, 93 (10), 1075–1075. https://doi.org/10.1094/pdis-93-10-1075b.

30. Galindo A.,J. Etiology of Planta Macho, a Viroid Disease of Tomato. Phytopathology 1982, 72 (1), 49. https://doi.org/10.1094/phyto-72-49.

31. Jin, J.; Tian, F.; Yang, D.-C.; Meng, Y.-Q.; Kong, L.; Luo, J.; Gao, G. PlantTFDB 4.0; Toward a Central Hub for Transcription Factors and Regulatory Interactions in Plants. Nucleic Acids Res 2016, 45 (Dl), Dl040–Dl045. https://doi.org/10.1093/nar/gkw982.

32. Zhou, M.; Memelink, J. Jasmonate-Responsive Transcription Factors Regulating Plant Secondary Metabolism. Biotechnol Adv 2016, 34 (4), 441–449. https://doi.org/10.1016/j.biotechadv.2016.02.004.(52)

33. Catinot, J.; Huang, J.-B.; Huang, P.-Y.; Tseng, M.-Y.; Chen, Y.-L.; Gu, S.-Y.; Lo, W.-S.; Wang, L.-C.; Chen, Y.-R.; Zimmerli, L. ETHYLENE RESPONSE FACTOR 96 Positively RegulatesArabidopsisresistance to Necrotrophic Pathogens by Direct Binding to GCC Elements of Jasmonate - and Ethylene-Responsive Defence Genes. Plant Cell Environ 2015, 38 (12), 2721–2734. https://doi.org/10.1111/pce.12583.

34. Feller, A.; Machemer, K.; Braun, E. L.; Grotewold, E. Evolutionary and Comparative Analysis of MYB and BHLH Plant Transcription Factors. Plant J 2011, 66 (1), 94–116. https://doi.org/10.1111/j.1365-313x.2010.04459.x.

35. Rehman, S.; Mahmood, T. Functional Role of DREB and ERF Transcription Factors: Regulating Stress-Responsive Network in Plants. Acta Physiol Plant 2015, 37 (9). https://doi.org/10.1007/sl1738-015-1929-1.

36. Gutterson, N. Regulation of Disease Resistance Pathways by AP2/ERF Transcription Factors. Curr Opin Plant Biol 2004. https://doi.org/10.1016/s1369-5266(04)00066-4.

37. Sacharowski, S.; Gratkowska, M.; Sarnowska, E.; et al. SWP73 Subunits of Arabidopsis SWI/SNF Chromatin Remodeling Complexes Play Distinct Roles in Leaf and Flower Development. Plant Cell 2015, 27(7), 1889–1906. https://doi.org:10.1105/tpc.15.00233

38. Yuan, L. Clustered ERF Transcription Factors: Not All Created Equal. Plant Cell Physiol 2020, 61 (6), 1025–1027. https://doi.org/10.1093/pep/pcaa067.

39. Wang, Y.; van der Hoeven, R.; Nielsen, R.; Mueller, L.; Tanksley, S. Characteristics of the tomato nuclear genome as determined by sequencing undermethylated EcoRI digested fragments. Theor Appl Genet 2005, 112(1) :72–84. https://doi.org/10.1007/s00122-005-0107-z

40. Peluso, J.; Delidow, B.; Lynch, J.; White, B. Follicle-stimulating hormone and insulin regulation of 17 beta-estradiol secretion and granulosa cell proliferation within immature rat ovaries maintained in perifusion culture. Endocrinology 1991, 128(1) :191–196. https://doi.org/10.1210/endo-128-1-191

41. Solano, R.; Stepanova, A.; Chao, Q.; Ecker, J. Nuclear events in ethylene signaling: a transcriptional cascade mediated by ETHYLE N E-I NSENSITIVE3 and ETHYLE N E-RESP ONSE-FACTORl. Genes Dev 1998, 12(23), 3703–3714. https://doi.org/10.l101/gad.12.23.3703

42. Staudt, A.; Wenkel, S. Regulation of Protein Function by “MicroProteins.” EMBO rep 2010, 12 (1), 35–42. https://doi.org/10.1038/embor.2010.196.

43. Eguen, T.; Straub, D.; Graeff, M.; Wenkel, S. MicroProteins: Small Size - Big Impact. Trends Plant Sci 2015, 20 (8), 477–482. https://doi.org/10.1016/j.tplants.2015.05.011.

44. Aviña-Padilla, K.; Ramírez-Rafael, J. A.; Herrera-Oropeza, G. E.; Muley, V. Y.; Valdivia, D. I.; Díaz-Valenzuela, E.; García-García, A.; Varela-Echavarría, A.; Hernández-Rosales, M. Evolutionary Perspective and Expression Analysis of Intronless Genes Highlight the Conservation of Their Regulatory Role. Front Genet 2021, 12. https://doi.org/10.3389/fgene.2021.654256.

45. Yang, C.; Huang, S.; Zeng, Y.; Liu, C.; Ma, Q.; Pruneda-Paz, J.; Kay, S. A.; Li, L. Two BHLH Transcription Factors, BHLH48 and BHLH60, Associate with Phytochrome Interacting Factor 7 to Regulate Hypocotyl Elongation in Arabidopsis. Cell Rep 2021, 35 (5), 109054. https://doi.org/10.1016/j.celrep.2021.109054.

46. Ahmad, Z. A Big Role for MicroProteins in Preventing Premature Floral Transition in the Shoot Meristem. Plant Physiol 2021, 187 (1), 12–13. https://doi.org/10.1093/plphys/kiab320.

47. Reverter, A.; Chan, E. K. F. Combining Partial Correlation and an Information Theory Approach to the Reversed Engineering of Gene Co-Expression Networks. Bioinformatics 2008, 24 (21), 2491–2497. https://doi.org/10.1093/bioinformatics/btn482.

48. Blondel, V. D.; Guillaume, J.-L.; Lambiotte, R.; Lefebvre, E. Fast Unfolding of Communities in Large Networks. J Stat Mech 2008, 2008 (10), P10008. https://doi.org/10.1088/1742-5468/2008/10/p10008.

49. Layat, E.; Cotterell, S.; Vaillant, I.; Yukawa, Y.; Tutois, S.; Tourmente, S. Transcript Levels, Alternative Splicing and Proteolytic Cleavage of TFIIIA Control 5S RRNA Accumulation during Arabidopsis Thaliana Development. Plant J 2012, 71 (1), 35–44. https://doi.org/10.1111/j.1365-313x.2012.04948.x.

50. Dissanayaka Mudiyanselage, S.; Qu, J.; Tian, N.; Jiang, J.; Wang, Y. Potato Spindle Tuber Viroid RNA-Templated Transcription: Factors and Regulation. Viruses 2018, 10 (9), 503. https://doi.org/10.3390/v10090503.

51. Liu, M.; Chen, Y.; Chen, Y.; Shin, J.-H.; Mila, I.; Audran, C.; Zouine, M.; Pirrello, J.; Bouzayen, M. The Tomato Ethylene Response Factor Sl-ERF.B3 Integrates Ethylene and Auxin Signaling via Direct Regulation of Sl-Aux/IAA27. New Phytol 2018, 219 (2), 631–640. https://doi.org/10.1111/nph.15165.

52. Srivastava, R.; Kumar, R. The Expanding Roles of APETALAZ/Ethylene Responsive Factors and Their Potential Applications in Crop Improvement. Brief Funct Genomics 2019, 18 (4), 240–254. https://doi.org/10.1093/bfgp/elz00l.

53. Fujimoto, S. Y.; Ohta, M.; Usui, A.; Shinshi, H.; Ohme-Takagi, M. Arabidopsis Ethylene-Responsive Element Binding Factors Act as Transcriptional Activators or Repressors of GCC Box-Mediated Gene Expression. Plant Cell 2000, 12 (3), 393. https://doi.org/10.2307/3870944.

54. Cheng, C. W.; Beech, D. J.; Wheatcroft, S. B. Advantages of CEMiTool for Gene Co-Expression Analysis of RNA-Seq Data. Comput Biol Med 2020, 125, 103975. https://doi.org/10.1016/j.compbiomed.2020.103975.

55. Aviña-Padilla K.;, Zambada-Moreno O.; Herrera-Oropeza GE.; Jimenez-Limas MA.; Hammond R.; Hudson M.; and Hernandez-Rosales*. Dynamic co-expression network analysis of root PSTVd-infected tomato reveals the interplay of BHLH TFs IntJ MoJ Sci in preparation

56. Abrahamiam, P. Analysis of SRNA Seq Data from TPMVd-Infected Tomato Plants, 2021.

57. Sukumari Nath, V.; Kumar Mishra, A.; Kumar, A.; Matoušek, J.; Jakše, J. Revisiting the Role of Transcription Factors in Coordinating the Defense Response against Citrus Bark Cracking Viroid Infection in Commercial Hop (Humulus Lupulus L.). Viruses 2019, 11 (5), 419. https://doi.org/10.3390/v11050419.

58. Márquez-Molins, J.; Villalba-Bermell, P.; Corell-Sierra, J.; Pallás, V.; Gómez, G. Integrative Time-Scale and Multi-Omic Analysis of Host-Responses to Hop Stunt Viroid Infection. 2022. https://doi.org/10.1101/2022.01.06.475203.

59. Gautier, L.; Cope, L.; Bolstad, B. M.; Irizarry, R. A. Affy--Analysis of Affymetrix GeneChip Data at the Probe Level. Bioinformatics 2004, 20 (3), 307–315. https://doi.org/10.1093/bioinformatics/btg405.

60. Scipy 08; 21, A.; Pasadena; Hagberg, A.; Swart, P.; Chult, D. S.; Los. Exploring Network Structure, Dynamics, and Function Using Networkx; 2008.

61. Kolberg, L.; Raudvere, U.; Kuzmin, I.; Vilo, J.; Peterson, H. Gprofiler2 -- an R Package for Gene List Functional Enrichment Analysis and Namespace Conversion Toolset G:Profiler. FlO00Res 2020, 9, 709. https://doi.org/10.12688/fl000research.24956.1.

62. Shannon, P. Cytoscape: A Software Environment for Integrated Models of Biomolecular Interaction Networks. Genome Res 2003, 13 (11), 2498–2504. https://doi.org/10.1101/gr.1239303.

63. Mercatelli, D.; Lopez-Garcia, G.; Giorgi, F. M. Corto: A Lightweight R Package for Gene Network Inference and Master Regulator Analysis. Bioinformatics 2020, 36 (12), 3916–3917. https://doi.org/10.1093/bioinformatics/btaa223

